# mTOR variants activation discovers PI3K-like cryptic pocket, expanding allosteric, mutant-selective inhibitor designs

**DOI:** 10.1101/2024.10.12.618044

**Authors:** Yonglan Liu, Wengang Zhang, Hyunbum Jang, Ruth Nussinov

## Abstract

mTOR plays a crucial role in PI3K/AKT/mTOR signaling. We hypothesized that mTOR activation mechanisms driving oncogenesis can advise effective therapeutic designs. To test this, we combined cancer genomic analysis with extensive molecular dynamics simulations of mTOR oncogenic variants. We observed that conformational changes within mTOR kinase domain are associated with multiple mutational activation events. The mutations disturb the α-packing formed by the kαAL, kα3, kα9, kα9b, and kα10 helices in the kinase domain creating cryptic pocket. Its opening correlates with opening of the catalytic cleft, including active site residues realignment, favoring catalysis. The cryptic pocket created by disrupted α-packing coincides with the allosteric pocket in PI3Kα can be harmoniously fitted by the PI3Kα allosteric inhibitor RLY-2608, suggesting that analogous drugs designed based on RLY-2608 can restore the packed α-structure, resulting in mTOR inactive conformation. Our results exemplify that knowledge of detailed kinase activation mechanisms can inform innovative allosteric inhibitor development.

## Introduction

Oncogenic mutations shift the conformational ensemble of their harboring oncoproteins (e.g., PI3K, K-Ras, B-Raf, ABL, SHP2, and mTOR)^1, 2, 3, 4, 5, 6, 7, 8, 9, 10, 11^ and tumor suppressors (e.g., p53, PTEN, and TSC1/2),^12, 13, 14^ increasing (decreasing) their catalytic capabilities. Hyperactivated signaling pathways promote cell growth, proliferation, and transformation, driving tumorigenesis.^15, 16, 17, 18^ Mutants and their corresponding wild types (WTs) often exhibit significant differences in sensitivity to the same regimen,^19, 20^ making mutation-specific therapeutics tailored to individual patients a significant clinical aim.^20, 21, 22, 23, 24, 25, 26^ Successful examples include the covalent sotorasib (AMG-510),^27^ which can effectively inhibit the K-Ras G12C mutant; the allosteric inhibitor RLY-2608,^19^ which selectively targets metastatic breast cancer with PI3Kα H1047R and E542K mutations; and the allosteric inhibitor STX-478,^20^ which selectively targets PI3Kα H1047R/L, E545K, and E542K mutations in advanced solid tumors and hormone receptor-positive (HR+) breast cancer. At the same time, patients harboring B-Raf V600E mutation are treated with the same drug regimen, regardless of cancer type and location.^28^ Sequencing technologies have advanced the detection of frequent tissue-specific genetic mutations, facilitating mutation-guided precision therapeutics.^29, 30, 31^ Mutations mapped onto protein structures guide drug development,^32^ underscoring the significance of structural and dynamic knowledge and insight into oncogenic activation of signaling proteins.^33, 34^

Mammalian target of rapamycin (mTOR) is a cornerstone of the PI3K/AKT/mTOR cell growth pathway, regulating diverse functions, including mRNA translation, thus cell size, autophagy, and metabolic processes.^35, 36, 37, 38^ In mammals, mTOR forms two core complexes, mTOR complex 1 (mTORC1) and complex 2 (mTORC2), predominantly activated at the lysosomal surface and plasma membrane, respectively.^39^ mTOR adopts a complex structure, permitting its evolving, intricate regulatory mechanisms. It primarily consists of the N-terminal HEAT (N-HEAT) repeats (residues 16-902), the middle HEAT (M-HEAT) repeats (residues 933-1241), the FAT (named after FRAP, ATM, and TRRAP) domain (residues 1261-2000), and the kinase domain (residues 2002-2549) (**Fig. S1a**).^40, 41, 42, 43, 44^ The kinase domain contains several inserted domains: the FKBP-rapamycin binding (FRB) domain (residues 2021-2118) in the N-lobe and the Lst8 binding element (LBE) domain (residues 2258-2296) and the FAT C-terminal (FATC) domain (residues 2517-2549) in the C-lobe.

Human cancer genomic databases point to mTOR mutations throughout its domains, with higher frequency observed in the FAT and kinase domains.^23, 45, 46^ mTORC1 includes the regulatory-associated protein of mTOR (Raptor), which facilitates substrate recognition of the TOR signaling motif in proteins such as ribosomal protein S6 kinase 1 (S6K1) and eukaryotic translation initiation factor 4E-binding protein 1 (4E-BP1).^47^ In mTORC2, mammalian stress-activated protein kinase interacting protein 1 (mSIN1) stimulates complex activation by anchoring to the membrane via its pleckstrin homology (PH) domain and aiding in substrate recruitment.^47, 48^ The mTOR’s FRB domain contains a secondary substrate-binding site that facilitates mTORC1 substrate recruitment.^44^ mTOR mutations that enhance substrate recruitment are expected to increase mTOR activity.^32^ The DEP domain-containing mTOR-interacting protein (DEPTOR) acts as an endogenous inhibitor of mTOR. Binding of DEPTOR to the mTOR FAT domain partially inhibits both mTORC1 and mTORC2.^49^ mTOR mutations that prevent the FAT-DEPTOR interaction could disrupt the balance between mTOR activation and inhibition, thereby enhancing mTOR signaling. GTP-bound Ras homolog enriched in brain protein (RHEB) allosterically activates mTORC1 by interacting with the N-HEAT. This interaction loosens the contact between the FAT and kinase domains (FAT KD), aligning active site residues for catalysis (**Fig. S1b**).^44^ Our previous studies, along with others, highlighted the critical role of the kα9b-helix (residues 2425-2436) and its overlapping region, the negative regulator domain (NRD, residues 2430-2450), in regulating mTOR activation.^50, 51, 52, 53, 54^ When these two motifs are positioned inside the catalytic cleft of mTOR, the cleft tends to adopt a deep and closed conformation. In contrast, positioning outside the catalytic cleft favors an open conformation, facilitating substrate entry. Mutations that promote an open catalytic cleft, disrupt FAT KD interactions, and align active site residues for catalysis enhance mTOR activity.

Here, we focus on mTOR, as there is still a significant lack of understanding of the structural and dynamic mechanisms underlying the oncogenic activation of its mutant forms. We seek to uncover commonalities and distinction in their activation mechanisms, which could assist drug interventions in genomics-guided and mutation-enriched “basket” trials.^23^ By comparing the structural and dynamic properties of the oncogenic activation mechanisms of variants, obtained from their extensive molecular dynamics (MD) simulations, with WT mTOR, we found that all mutations inherently promote mTOR activation by disrupting (or disturbing) the packing structure formed by helices kαAL, kα3, kα9, and kα10 centered around helix kα9b in the C-lobe of the kinase domain. Especially, this disruption correlates with the opening of the catalytic cleft, and the realignment of active site residues favorable for catalysis. Our findings further underscore the pivotal role of the FAT domain in the regulation of mTOR activity and decipher exactly how it conducts the regulation.

Especially, PI3K and mTOR are mechanistically similar. mTOR belongs to the PI3K-related kinases (PIKK) superfamily, with mTOR considered a part of the PI3K family due to its catalytic domain similarity to lipid kinases like PI3K.^55^ Even though overlooked to date, likely since it was not captured in crystal structures, it is not surprising that the mTOR mutant-promoted activation mechanisms that we determined discovered a cryptic pocket resembling that found in the C-lobe of the PI3Kα kinase domain.^19, 20, 56^ Molecular docking of the PI3Kα allosteric inhibitor RLY-2608 to the newly discovered cryptic pocket in mutant mTOR (I2500F) and the corresponding position in WT mTOR, followed by MD simulations of the resulting complexes, suggest that structure-based design of new mTOR allosteric inhibitors could selectively target mutant mTOR over WT mTOR. This cryptic pocket finding and inhibitor fitting, hold promise for the design of mutant-sensitive allosteric inhibitors of mTOR *by learning* from the allosteric inhibitors of PI3Kα. Encouragingly, just now Kevan Shokat and his collaborators observed a conserved, targetable cryptic pocket in members of the Ras, Rho, and Rab family of GTPases.^57^ While their pocket details differ, the authors underscore the potential that the pockets offer for the development of optimized inhibitors, as we have done here. Homologous cryptic pockets in same-family proteins are a powerful drug discovery concept.

## RESULTS

### mTOR mutational analysis from cancer genomic databases

To identify cancer-derived mutations in mTOR, we integrated data from two cancer genome databases, The Cancer Genome Atlas (TCGA, the MC3 MAF file)^58^ and Genomics Evidence Neoplasia Information Exchange (GENIE, v13.1-public), and conducted comprehensive analyses. After pre-filtering, we obtained 1447 missense mutations from 1373 samples across 1294 patients (**Fig. 1a** and **Supplementary File**). 199 different missense mutations of mTOR were identified, spanning 55 known cancer types, each occurring at varying frequencies and positions. These mutations were observed in 29 different tissues, with the highest occurrence in bowel, uterus, and kidney (**Fig. 1b**). In line with prior work,^23, 35^ most cancer-derived mutations occur within the FAT and kinase domains (∼70%) (**Fig. 1c**). The higher PolyPhen scores^59^ for mutant mTORs with mutations in the FAT and kinase domains than those with mutations in the HEAT regions indicate that mTOR mutations in the FAT and kinase domains generally have a greater impact on the protein conformation than mutations located in the N-HEAT and M-HEAT regions (**Fig. 1d**). Prominent mutation hotspots include residues C1483, H1647, E1799, T1977, V2006, S2215, E2419, L2427, and I2500 (**Fig. 1e**). Activating mutations linked to cancer rely primarily on residues that stabilize active protein conformations.^60, 61^ Certain hotspots, such as C1483 and H1647, are located at the intra-FAT hinge, distant from the kinase domain (**Fig. 1f**). Others, such as residues E1799 and E2419, reside at the FAT–α-packing interface. T1977 and V2006 are clustered within the anchoring region linking the C-terminal region of the FAT domain (FAT^C-term^) to the N-lobe of the kinase domain (KD^N-lobe^). Mutations at L2427 and I2500 occur within the α-packing regions. S2215 is positioned at the kα3b-helix in the N-lobe, presenting a challenge to predict the impact of mutations at this site on the conformation of mTOR.

**Fig. 1.**
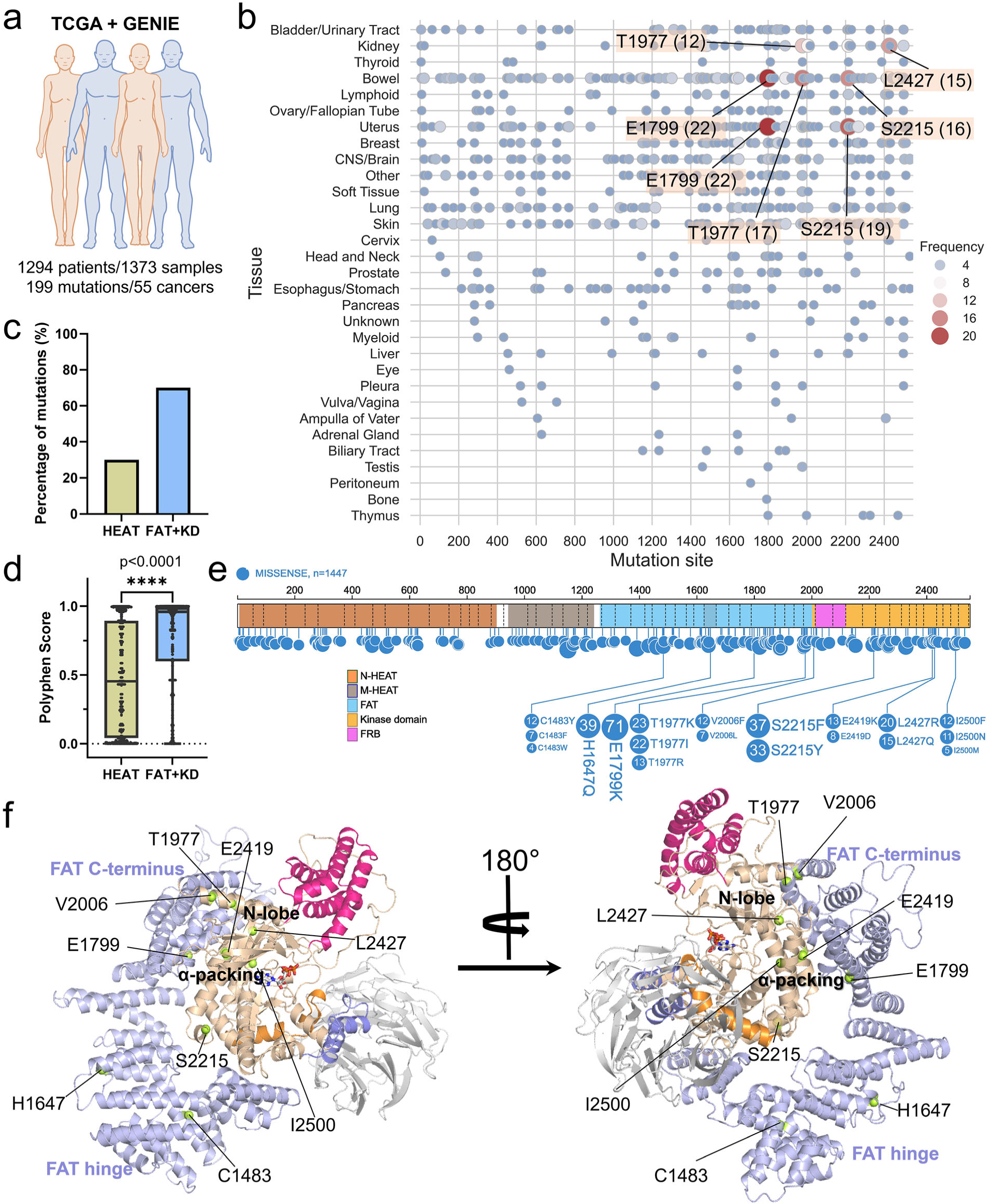
Identification of cancer-derived mutations of mTOR. (a) Overview of cancer-derived mutations of mTOR, detailing the number of patients, samples, mutations, and associated cancers sourced from TCGA and GENIE cancer genomic databases. (b) Mutation frequencies of mTOR at the tissue level. (c) Percentages of mTOR mutations occurring in the HEAT (N-HEAT and M-HEAT) and FAT+KD (the FAT and kinase domains) regions. (d) PolyPhen scores of mTOR mutations in the two regions. PolyPhen scores range from 0.0 (tolerated) to 1.0 (deleterious), indicating the predicted impact of mutations. Scores closer to 1.0 suggest higher potential for deleterious effects. A nonparametric Mann-Whitney test (p < 0.0001) on PolyPhen scores reveals that mutations within the FAT-KD region induce significantly higher deleterious effects compared to those occurring in the HEAT region. (e) Mutation frequencies of mTOR at the residue level. (f) Mapping of hot-spot cancer mutation sites onto the mTOR structure.

### FAT domain regulation of mTOR

Physiologically, RHEB allosterically activates mTORC1, initiated by its physical interaction with the N-HEAT of mTOR.^44, 62^ This loosens the contact between the FAT and kinase domains of mTOR, allowing the active site residues to adopt orientations favorable for catalysis. These scenarios suggest a critical role for the FAT domain in the regulation of mTOR kinase activity. Here, we provide atomic and dynamic data to reveal how this domain is involved in regulation and shed light on its subsequent impact on mTOR activity. We generated two simulation models based on the crystal structure (PDB ID: 4JSP), which is a complex consisting of the N-terminal truncated mTOR (residues 1382-2549) and mLST8. This complex exhibits kinase activity comparable to that of mTORC1 (**Fig. S2 and Table S1**).^50^ One model is the complex of mTOR with the FAT domain deleted (system name: ΔFAT), i.e. only the kinase domain, and the other is the WT complex with the FAT domain (system name: WT).

During the simulations, we observed a notable difference in the conformation of the mTOR kinase domain between the ΔFAT and WT systems. Principal component analysis (PCA) reveals the distinct clustering patterns formed by the first two PCs for the kinase domain, indicating that deletion of the FAT domain causes a shift in the conformational ensemble of this domain (**Fig. 2a**). The activation loop (A-loop, residues 2357-2379) of mTOR contains an inserted kαAL-helix motif (residues 2363-2367) that participates, with three other helices (kα3, kα9, and kα10), in the packing centered around the kα9b-helix in the C-lobe of the kinase domain. We refer to this arrangement as the kαAL-kα3-kα9-kα9b-kα10 packing (or α-packing). In the WT system, the α-helical conformation of the kαAL-helix and the α-packing are well maintained (**Fig. 2b and 2c**). However, in the absence of the FAT domain, the kαAL-helix completely loses its helical structure, resulting in an extended and highly fluctuating A-loop, in contrast to the collapsed and stable A-loop observed in the WT system (**Fig. 2b**). This is evident from the greater distance from the ATP and larger root-mean-square-fluctuations (RMSFs) of A-loop for the ΔFAT system than those for the WT system (**Fig. 2d and 2e**). The catalytic loop (Cat-loop, residues 2336-2343) contains a negatively charged residue, D2338, which is required for mTOR phosphotransferase activity. mTOR with the D2338A mutation is a well-established kinase-dead variant, known for its capacity to abolish phosphotransferase activity and inhibit mTOR signaling both *in vitro* and *in vivo*.^50, 63, 64, 65, 66, 67, 68^ When D2338 is positioned in close proximity and properly oriented toward ATP, it is anticipated to facilitate efficient transfer of the phosphate group from ATP to the substrate; if D2338 is distant, it may reduce the effectiveness of transferring.^3, 44^ In the collapsed A-loop, the catalytic residue D2338 forms a salt bridge with the positively charged residue R2368 in the A-loop, which hinders its orientation and proximity to ATP (**Fig. 2f***, top left*). However, in the ΔFAT system, with the A-loop extension, the D2338-R2368 salt bridge has never occurred (**Fig. 2f***, top right*). As a result, the catalytic residue D2338 approaches and orients toward ATP, potentially enhancing catalytic efficiency, as indicated by the shorter *d_ATP-D2338_* and longer *d_D2338-R2368_* for the ΔFAT system compared to the WT system (**Fig. 2f***, bottom panels*). Apart from D2338, three other residues—H2440 and N2343 in the Cat-loop, and D2357 in the A-loop—are crucial for catalysis as well.^44, 50^ H2440 serves as another catalytic residue, and N2343 and D2357 function as metal ligand binders, coordinating magnesium ions. During the simulations, we did not observe discernible differences in these three residues between these two systems. The conformational change in the A-loop of mTOR has the potential to affect the kα9b-helix. Although we did not observe a significant “OUT” motion of the kα9b-helix in the ΔFAT system within the simulation time, the visualization shows a loss of some helical components (**Fig. 2b**). In the ΔFAT system, both the kα9b-helix and the NRD exhibit higher fluctuations compared to the WT system (**Fig. 2g**), indicating instability within the catalytic cleft. Taken together, these contrasting behaviors highlight the critical regulatory role of the FAT domain in mTOR activity. Stable and proper FAT-KD contact inhibits mTOR by stabilizing the α-packing. This stabilization partially obstructs the catalytic cleft and induces active site residues to adopt orientations less favorable for catalysis.

**Fig. 2.**
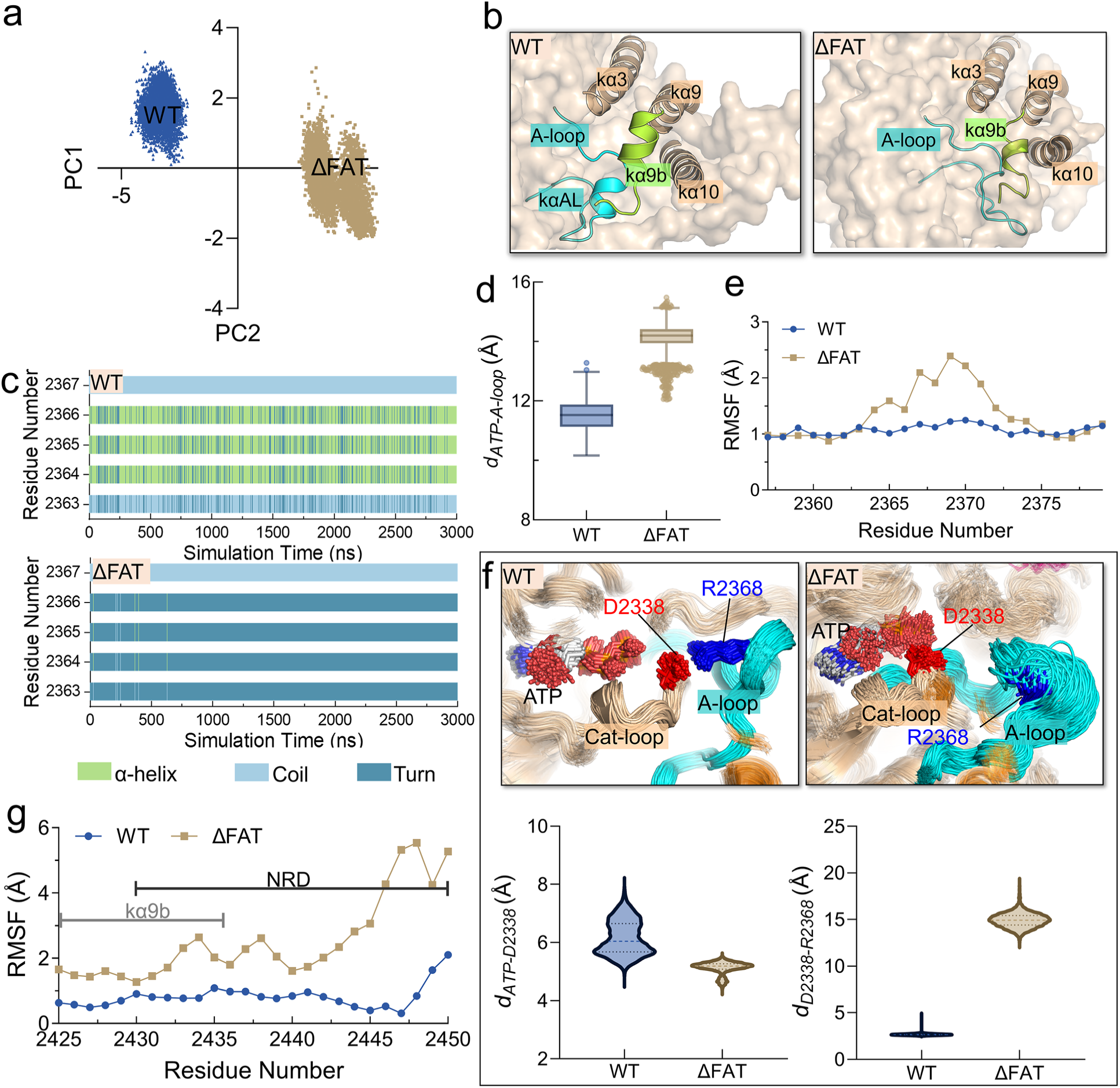
FAT domain regulation of mTOR activity. (a) Projections of PC1 and PC2 from the PCA for the kinase domain of mTOR, (b) structural representations of the packing structures formed by the kαAL, kα3, kα9, kα9b, and kα10 helices, (c) time-dependent (0-3 μs) of the secondary structures of the kαAL-helix (residues 2363-2367), (d) boxplots depicting *d_ATP_*_□*A-loop*_ [distance between ATP:PG and the center of mass of Cα atoms of partial A-loop region (residues 2357-2380)], (e) RMSFs of A-loop, (f) conformational ensembles of the A-loop, R2368 in the A-loop, D2338 in the Cat-loop, and the ATP and violin plots illustrating *d_ATP_*_□*D2338*_ (distance between ATP:PG and D2338:OD2) and *d_D2338_*_□*R2368*_ (distance between D2338:OD2 and R2368:NH1), (g) RMSFs of the kα9b-helix and the NRD for the WT and ΔFAT systems. The secondary structures were calculated using the DSSP plugin in VMD.

### Oncogenic activation of mTOR variants

The activation principles of mutant variants of mTOR can be either similar or different. Understanding these oncogenic activation mechanisms at the atomic level is crucial for developing innovative effective allosteric therapeutic strategies against cancer driven by mTOR mutations. This could potentially enhance mutation-enriched "basket" trials in precision medicine.^21, 23^ To delve deeper into these mechanisms, we chose several hotspot mutants— E1799K, T1977K, V2006F, S2215F, E2419K, L2427R, and I2500F—for MD simulations.

### T1977K, V2006F, and S2215F mutations, which disrupt the anchoring of the FAT^C-term^ to the KD^N-lobe^, opening the catalytic cleft and increasing substrate accessibility

The FAT^C-term^ is tightly anchored to the KD^N-lobe^ of mTOR through the interaction between the fα33-helix of the FAT domain and the kα1-helix and kβ6-strand (residues 2179-2187) of the kinase domain (**Fig. S3**). Immediately adjacent to the FRB domain is the kα1-helix, which appears to be responsible for anchoring the FRB domain to the N-lobe of the kinase domain. We hypothesized that mutations that disrupt the interaction of the fα33-helix with the N-lobe could alter the dynamics of the FRB domain. The catalytic cleft of mTOR adopts a deep "V-shape" conformation, characterized by mLST8 and the FRB domain of mTOR extending from the C-lobe and the N-lobe, respectively. This unique structure limits substrate access to the ATP-binding site. Experimental data showed that deletion of NRD increased mTOR activity by 3.5 fold.^53^ Observations from MD simulations reveal that the mTOR/mLST8 complex lacking NRD (system name: ΔNRD) prefers an open and shallow catalytic cleft, which favors substrate access (**Fig. 3a and 3b**). This provides an atomic-level explanation for why removal of NRD was able to enhance mTOR’s activity.^51^ In our simulations, we observed a notable conformational shift in the "V-shaped" catalytic cleft with the T1977K, V2006F, and S2215F mutations, transitioning from a closed to an open state (**Fig 3b**). This transition is evident from the significantly longer *d_FRB-LBE_* observed for these mutants (∼41 Å for T1977K, ∼33 Å for V2006F, and ∼34 Å for S2215F) compared to the WT mTOR (∼23 Å) (**Fig. 3a**). This suggests that the T1977K, V2006F, and S2215F mutations may enhance the accessibility of the catalytic cleft, potentially facilitating substrate entry to reach the ATP binding site. The T1977K and V2006F mutations, located in the fα34-helix and kα1-helix, respectively (**Fig. S4 and Table S1**), are likely to disrupt the anchoring of the FAT^C-term^ to the KD^N-lobe^, thereby increasing the dynamics of the FRB domain. Although S2215 is located in the kα3b-helix away from the FAT^C-term^ KD^N-lobe^ interface, its mutation to phenylalanine (S2215F) alters the conformation of kβ6-strand (**Fig. S5**), which can allosterically affect the interaction with the fα33-helix, resulting in an effect similar to that observed with the T1977K and V2006F mutations.

**Fig. 3.**
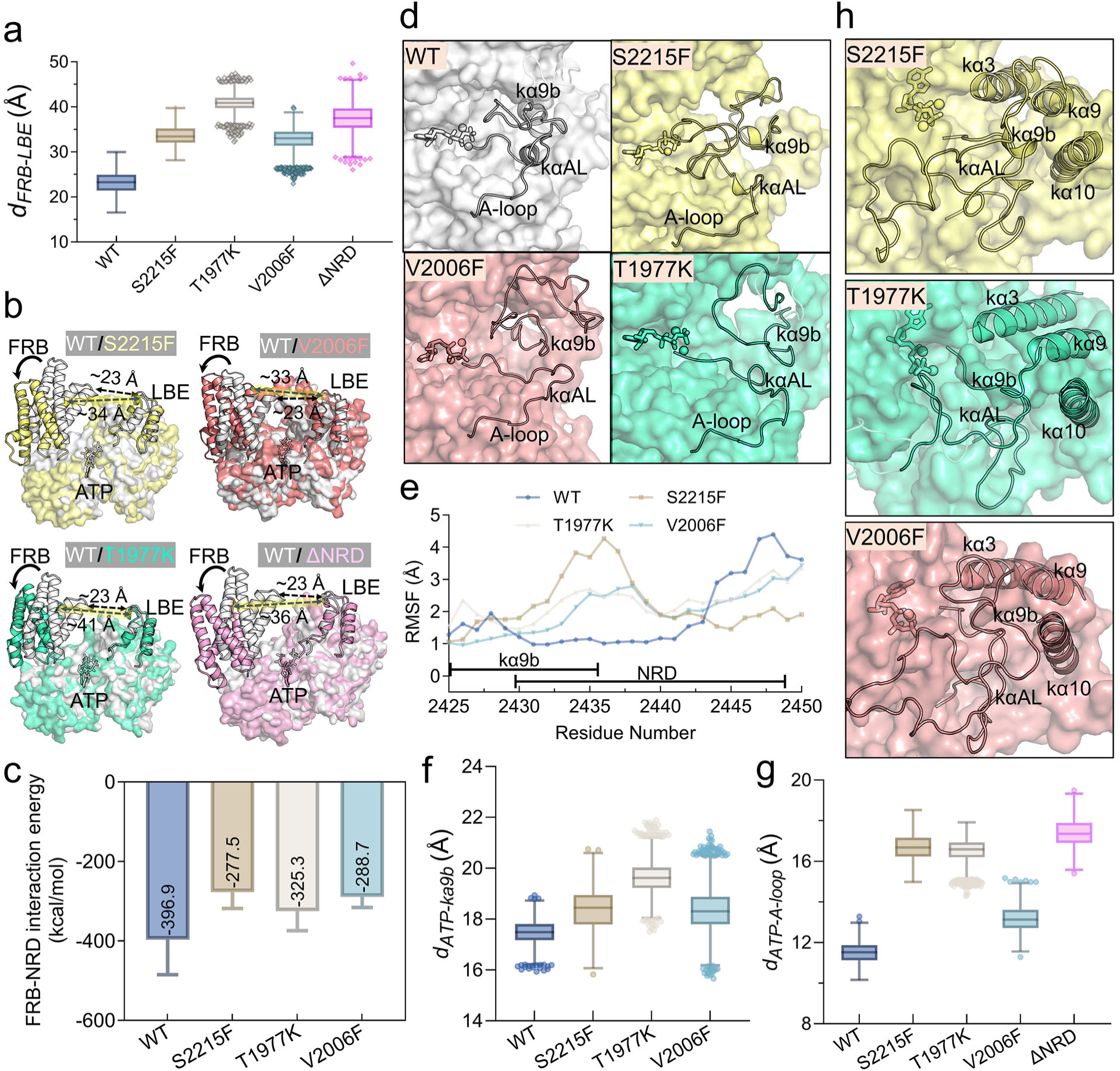
Mutations disrupting the anchoring of FAT’s C-terminal region onto kinase domain’s N-lobe. (a) Boxplots illustrating *d_FRB-LBE_* (distance between the Cα atom of tM2039 in the FRB domain and the Cα atom of M2277 in the LBE domain) for the WT, S2215F, T1977K, V2006F, and ΔNRD systems. (b) Structural alignment comparing the catalytic cleft of the WT system with the S2215F, T1977K, V2006F, and ΔNRD systems. (c) Interaction energies between the NRD and the FRB domain, (d) representative snapshots depicting the conformation of kαAL, the A-loop, and kα9b, (e) RMSFs of kα9b and the NRD, and boxplots depicting (f) *d_ATP_*_□*k*α_*_9b_* and (g) *d_ATP_*_□*A-loop*_ for the WT, S2215F, T1977K, V2006F, (and NRD) systems. (g) Representative snapshots showing the packing structures formed by the kαAL, kα3, kα9, kα9b, and kα10 helices for the three mutant systems (S2215F, T1977K, and V2006F).

In the WT, the NRD, located in the catalytic cleft, acts as a glue that restricts the movement of FRB. This restriction is facilitated by the positively charged residues (K2440, R2441, R2443) of the NRD, which form salt bridges and electrostatic interactions with the negatively charged residues (E2032 and E2033) in the FRB helix (**Fig. S6**).^51^ The movement and displacement of the FRB domain weaken its confinement by the NRD, as indicated by a significant increase in the interaction energy in these three mutants compared to the WT system (**Fig. 3c**). In stark contrast to the stable α-helical conformation (**Fig. 3d**) and lower RMSFs (**Fig. 3e**) observed in the kα9b-helix of WT mTOR, the T1977K, V2006F, and S2215F mutants exhibit a coil structure and higher RMSFs in this motif. These changes in the mutants potentially lead to an outward movement of the kα9b-helix (**Fig. 3f**), as evidenced by their longer separation distances (∼18.5/∼19.8/∼18.4 Å for S2215F/T1977K/V2006F) between ATP and kα9b with respect to that of WT mTOR (∼17.4 Å). Similar to mTOR lacking the FAT domain, the T1977K, V2006F, and S2215F mutants undergo a conformational change from α-helix to coil or turn structures in the kαAL-helix (**Fig. S7**), along with an extension of the A-loop (**Fig. 3d and 3g**). These changes are accompanied by the disruption of the α-packing (**Fig. 3h**), supporting previous proposals.^44, 50, 51^ However, for the catalytic residue D2338 in these three mutants, no preference for orientation toward ATP is observed within the simulation time (**Fig. S8**). In summary, the T1977K, V2006F, and S2215F mutations can lead to an open catalytic cleft, outward movement of the kα9b-helix, extension of the A-loop, and disruption of α-helix packing, thereby enhancing substrate accessibility to the catalytic cleft.

### E1799K and E2419K mutations weaken FAT–**α**-packing interface

The E1799K and E2419K mutations occur at the FAT α-packing interface (**Fig. 1f**). In WT mTOR, there are two stable salt bridges at this interface: one is between E1799 in the fα28-helix of the FAT domain and R2505 in the kα10-helix of the kinase domain (E1799-R2505), and the other is between E2419 in the kα9-helix of the kinase domain and R1905 in the fα30-helix of the FAT domain (E2419-R1905) (**Fig. 4a**). The E1799K substitution directly disrupts the E1799-R2505 salt bridge but does not affect the E2419-R1905 salt bridge. The E2419K mutation disrupts both the E2419-R1905 and E1799-R2505 salt bridges. Both mutations result in a looser FAT–α-packing contact (**Fig. 4b**). We hypothesized that the E1799K and E2419K mutations may induce conformational changes in the kinase domain resembling those observed with FAT deletion to promote mTOR activation. As expected, like the ΔFAT system, the two mutants fail to maintain the α-helical structure of the kαAL-helix in residues 2364-2366, which remains stable in the WT system (**Fig. 4c**). In the E1799K mutant, the conformational change of the A-loop coincides with the disruption of the α-packing. Of particular note is the complete loss of the α-helical structure in the kα9b-helix (**Fig. 4d**). While the kα9b helix maintains its conformation in the E2419K mutant, its dynamics are significantly heightened compared to the WT system. The loss of the α-helical structure in kα9b in the E1799K mutant and the increased dynamics in the E2419K mutant reduce the interaction between the FRB domain and the NRD, potentially diminishing the restriction on the FRB domain (**Fig. 4e and 4f**). This may increase the likelihood that the kα9b-helix and the NRD shift away from the catalytic cleft, thereby opening it and making the active site more accessible to substrates. The A-loop also tends to extend in both mutants, but less so than in the ΔFAT system (**Fig. 4d and 4g**). The extension of the A-loop also disrupts the D2338-R2368 salt bridge in both systems. D2338 near ATP is observed in the E1799K mutant but not in the E2419K mutant (**Fig. S9**), indicating that the proximity of D2338 to ATP in the E2419K mutant may occur over a longer timescale. Nevertheless, we observed that D2338 has a greater tendency to orient to ATP in both mutants compared to the WT system (**Fig. 4h**). This orientation can potentially increase their phosphotransferase activity. Overall, the E1799K and E2419K mutations weaken the interaction between FAT and the N-lobe, leading to disruption of the α-helix packing and causing catalytic residues to adopt orientations conducive to catalysis.

**Fig. 4.**
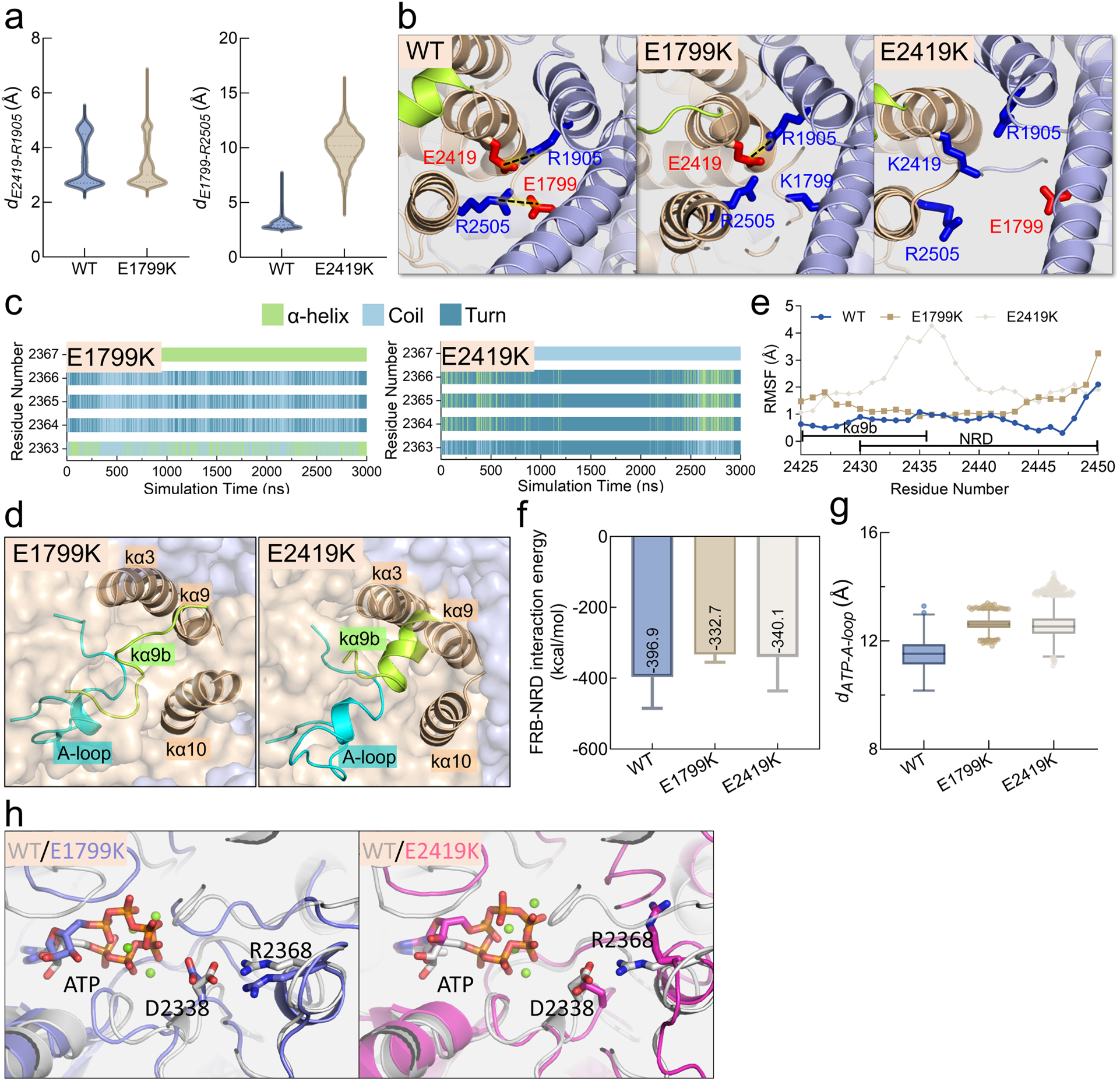
Mutations weakening the FAT–α-packing interface. (a) Volin plots depicting *d_E2419-R1905_* (distance between E2419:OE2 and R1905:NH1, *left*) for the WT and E1799K systems and *d_E1799-R2505_* (distance between E1799:OE2 and R2505:NH1, *right*). (b) Representative snapshots showing the interface between the FAT domain and the α-packing of the kinase domain with salt bridges indicated by dashed lines, (c) time-dependent (0-3 μs) of the secondary structures of kαAL, (d) RMSFs of kα9b and the NRD, (e) representative snapshots displaying the α-packing, (f) interaction energies between the NRD and the FRB domain, (g) boxplots of *d_ATP_*_□*A-loop*_, and (h) representative snapshots showing the orientations of residues D2338 and R2468 for the WT, E1799K, and E2419K systems.

### L2427R and I2500F mutations directly disrupt **α**-packing

L2427 and I2500 are located in the α-packing region, with L2427 located in the kα9b-helix and I2500 in the kα10-helix. In WT mTOR, the side chains of these two residues are oriented inwards within the α-packing, whereas both mutations directly disrupt this packing (**Fig. 5a**). Throughout the simulations, the I2500F mutation leads to the loss of the α-helical structure in residues 2364-2366 of the kαAL motif (**Fig. 5b**). Although the L2427R mutant retains most of the α-helical structure in this motif, it is not as stable as observed in WT mTOR. Compared to WT mTOR, the significant increase in *d_ATP-A-loop_* suggests that the A-loop in the L2427R and I2500F mutants has shifted further away from the catalytic site (**Fig. 5c**), like other mutant mTORs and mTOR lacking the FAT domain (**Fig. 3g and 4g**). In particular, it is fully extended in the I2500F mutant (**Fig. 5a and 5c**). Consistent with our previous studies,^51^ the helical structure of the kα9b completely disappears, resulting in higher fluctuations. Although the I2500F mutation maintains the helical component of kα9b, the dynamics of this motif increase significantly (**Fig. 5d**), suggesting instability in its position or conformation. The conformational change and heightened dynamics of the kα9b-helix can directly affect the dynamics of the NRD. This reduces the constraint of the NRD on the FRB domain, as evidenced by the notable increase in the interaction energy between the FRB domain and the NRD for these two mutants (∼-142.6/∼-218.5 kcal/mol for L2427R/I2500F) comparing to WT mTOR (**Fig. 5e**). During the simulations, we did not observe a significant tendency for the catalytic residue D2338 to approach or orient toward ATP (**Fig. S10**), suggesting that longer timescales may be required to observe such behavior. Taken together, the L2427R and I2500F mutations directly disrupt the α-packing, reducing the constraint of the kα9b-helix on the FRB domain. This change can potentially lead to rotation of the FRB domain, resulting in the opening of the catalytic cleft.

**Fig. 5.**
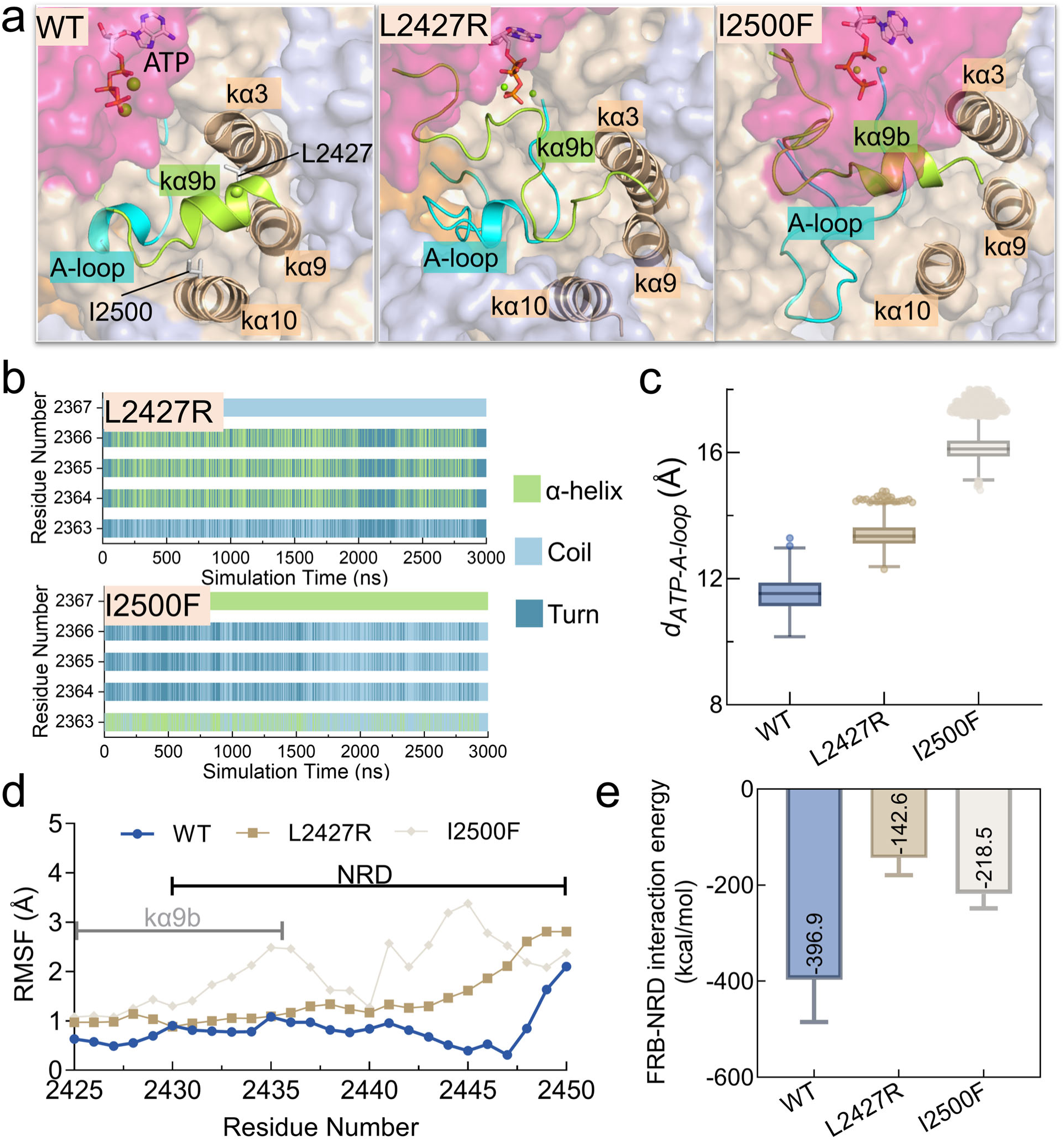
Mutations directly disrupting the α-packing. (a) Representative snapshots displaying the α-packing, (b) time-dependent (0-3 μs) of the secondary structures of the kαAL-helix, (c) boxplots of *d_ATP_*_□*A-loop*_, (d) RMSFs of kα9b and the NRD, (e) interaction energies between the NRD and the FRB domain for the WT, L2427R, and I2500F systems.

### PI3K**α** inhibitors, and mTOR activation mechanisms, inform mutant-selective allosteric inhibitor development

mTOR is a critical drug target with three generations of inhibitors: allosteric (rapamycin and rapalogs), orthosteric, and bitopic inhibitors.^69^ Clinical research has led to the approval of rapalogs such as everolimus and temsirolimus for cancer treatment. While effective against WT mTOR, these inhibitors are not selective for mutants. Our simulations of mutant mTORs reveal that all selected mutations can disrupt the α-packing in the kinase domain, shifting the conformational ensemble toward configurations that favor substrate access to ATP or optimize the alignment of active site residues for efficient phosphoryl transfer during catalysis. Stabilizing the α-packing can potentially inhibit mutant mTORs, suggesting a strategy for developing mutant-selective inhibitors.

In WT mTOR, the kαAL-helix and F2362 in the collapsed region of the A-loop dock into a pocket formed by helices kα3, kα6, kα9, kα9b, and kα10 (**Fig. 6a**). Here, we used the I2500F mutant as a case study because of its significantly extended A-loop. In the I2500F mutant, F2362 forms π-π stacking with F2500 (**Fig. S11**), shifting the A-loop away from the catalytic cleft and creating a cryptic allosteric pocket originally occupied by F2362 (**Fig. 6b**). mTOR shares structural similarities with the PI3K family in the kinase domain. Mutant-selective allosteric inhibitors, such as STX-478 and RLY-2608,^19, 20, 70^ bind to PI3K at a site identical to the newly identified cryptic pocket in the I2500F mutant mTOR (**Fig. 6c**). In p110α of PI3Kα, F397 reflects the orientation of F2362 in WT mTOR. Upon binding of RLY-2608, this position is occupied by the 2-chloro-5-fluorophenyl group of RLY-2608 and induces the α-helix conformation of the A-loop. We aligned the cryptic pocket of the I2500F mutant mTOR with the RLY-2608 bound to PI3Kα and observed that RLY-2608 fits well into this pocket, with its 2-chloro-5-fluorophenyl group occupying the original position of F2362, albeit with slight overlap with several residues in the A-loop (**Fig. 6d**). Notably, drug binding can promote local conformational changes that create significant geometric features within a binding site.^34, 71, 72^ RLY-2608 shows significantly greater selectivity for the H1047R PI3Kα mutant (IC_50_ = 4 ± 0 nmol/L) compared to WT PI3Kα (IC_50_ = 48 ± 17 nmol/L), primarily because the mutant kinase domain is less constrained by certain residues and motifs within or near the binding pocket.^70^ The disrupted and more perturbed α-packing in the mutant mTOR reduces the energy barrier and renders higher stability of the allosteric drug binding to the cryptic pocket compared to the WT, leading to higher drug residence time. This difference provides the rationale for the selectivity of the mutant mTORs over the WT form.

**Fig. 6.**
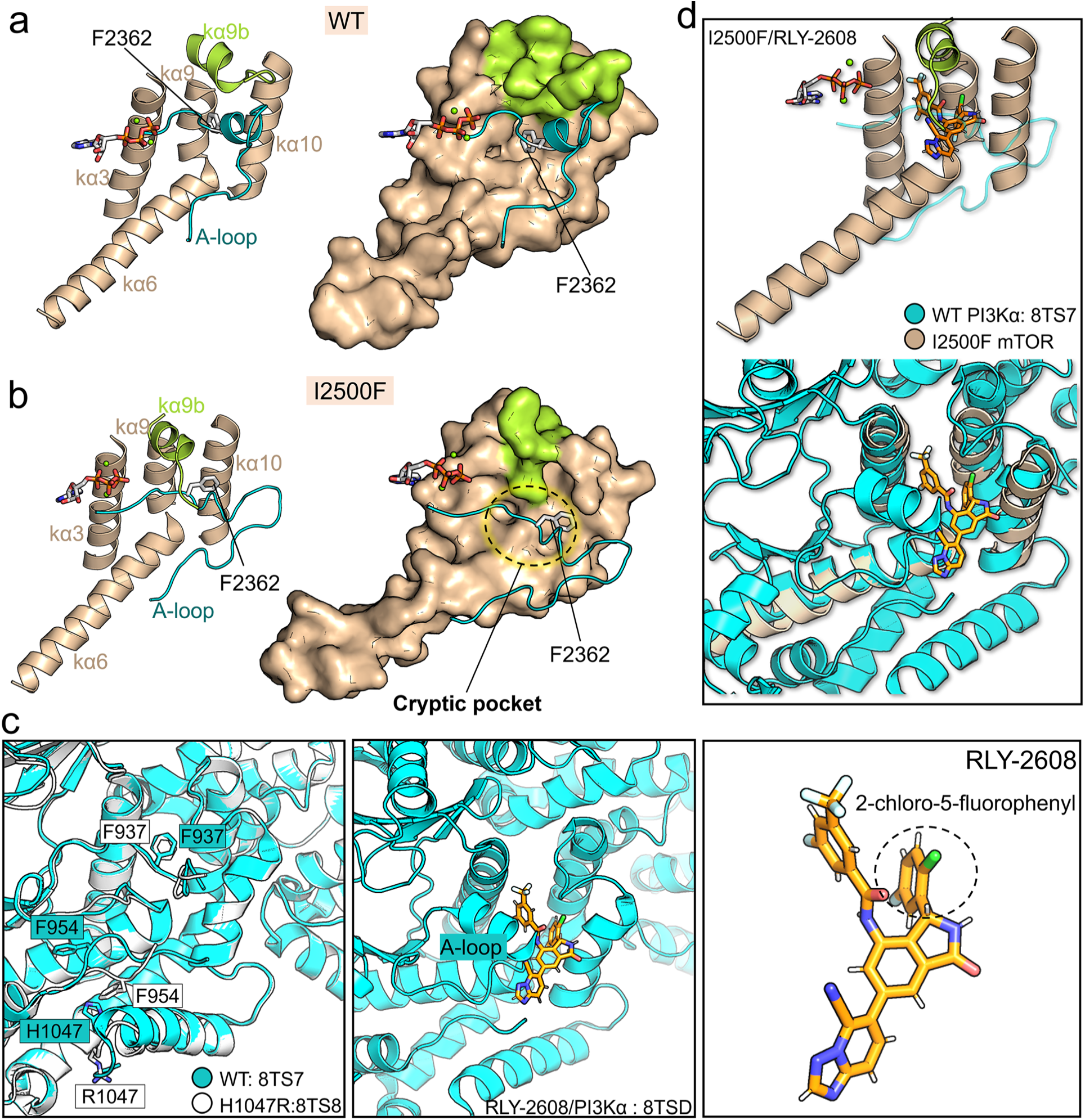
Mutant-selective cryptic pocket of mTOR. Orientation of residue F2362 and the pocket formed by kα3, kα6, kα9, kα9b, and kα10 helices in (a) the WT and (b) I2500F systems. (c) *Left*: orientation of residue F937 in the unbounded state of WT (PDB ID: 8TS7) and H1047R mutant (PDB ID: 8TS8) forms of PI3Kα; *middle*: binding posture of allosteric inhibitor RLY-2608 at the cryptic pocket of p110α subunit of PI3Kα (PDB ID: 8TSD); *right*: chemical structure of RLY-2608. (d) *Top*: modeling of the cryptic pocket found in the I2500F mutant mTOR bound with RLY-2608; *below*: structural alignment of this cryptic pocket with the one in the p110α subunit of PI3Kα.

To inform the design of an allosteric mTOR inhibitor based on the structure of RLY-2608, we conducted induced fit docking (IFD) followed by MD simulations and compared the binding of RLY-2608 to WT mTOR, the I2500F mutant mTOR, and to PI3Kα. Throughout the simulations we observed that RLY-2608 maintains stable binding to all three proteins. As expected, the 2-chloro-5-fluorophenyl group in RLY-2608 stably occupies the position of F2362 of the WT and mutant mTORs. Notably, RLY-2608 exhibits more favorable binding to the I2500F mutant than to WT mTOR, and its affinity for the mutant is comparable to that to PI3Kα (**Fig. 7a**). This suggests that RLY-2608 has a greater selectivity for the I2500F mutant than for the WT mTOR. RLY-2608 binding induces a conformational change in the A-loop of the I2500F mutant mTOR, shifting it from an extended conformation to a collapsed one, accompanied by the formation of an α-helix (**Fig. 7b**). Upon binding, RLY-2608 forms three hydrogen bonds (H-bonds) with N2435, C2362, and L2334, as well as hydrophobic interactions with V2504, V2422, F2421, L2418, F2202, L2201, and V1198. N2435, located in the NRD, adopts a favorable orientation upon drug binding, enabling the formation of an additional hydrogen bond with D2338 (**Fig. 7c & 7d**). This interaction likely prevents N2435 from orienting towards the ATP-binding site, potentially inhibiting efficient catalysis (**Fig. 7e**). However, RLY-2608 consistently displays bad interactions with F2500 due to steric hindrance between the bulky ring of the γ-lactam group in RLY-2608 and F2500 of mTOR (see purple dashed lines in **Fig. 7d**). To improve the selectivity for the cryptic pocket of the I2500F mutant mTOR, we propose to modify the inhibitor’s structure by reducing the size of the 5-membered ring in this group. However, since other mutant mTORs do not contain F2500, this modification may not be applicable to them. Alternatively, a cysteine residue (C2361) in the A-loop is close to the drug binding pocket, providing an opportunity for covalent drug design (**Fig 7f**). Targeting C2361 for covalent attachment could enhance the binding affinity and stability of the newly designed drugs.^73, 74^

**Fig. 7.**
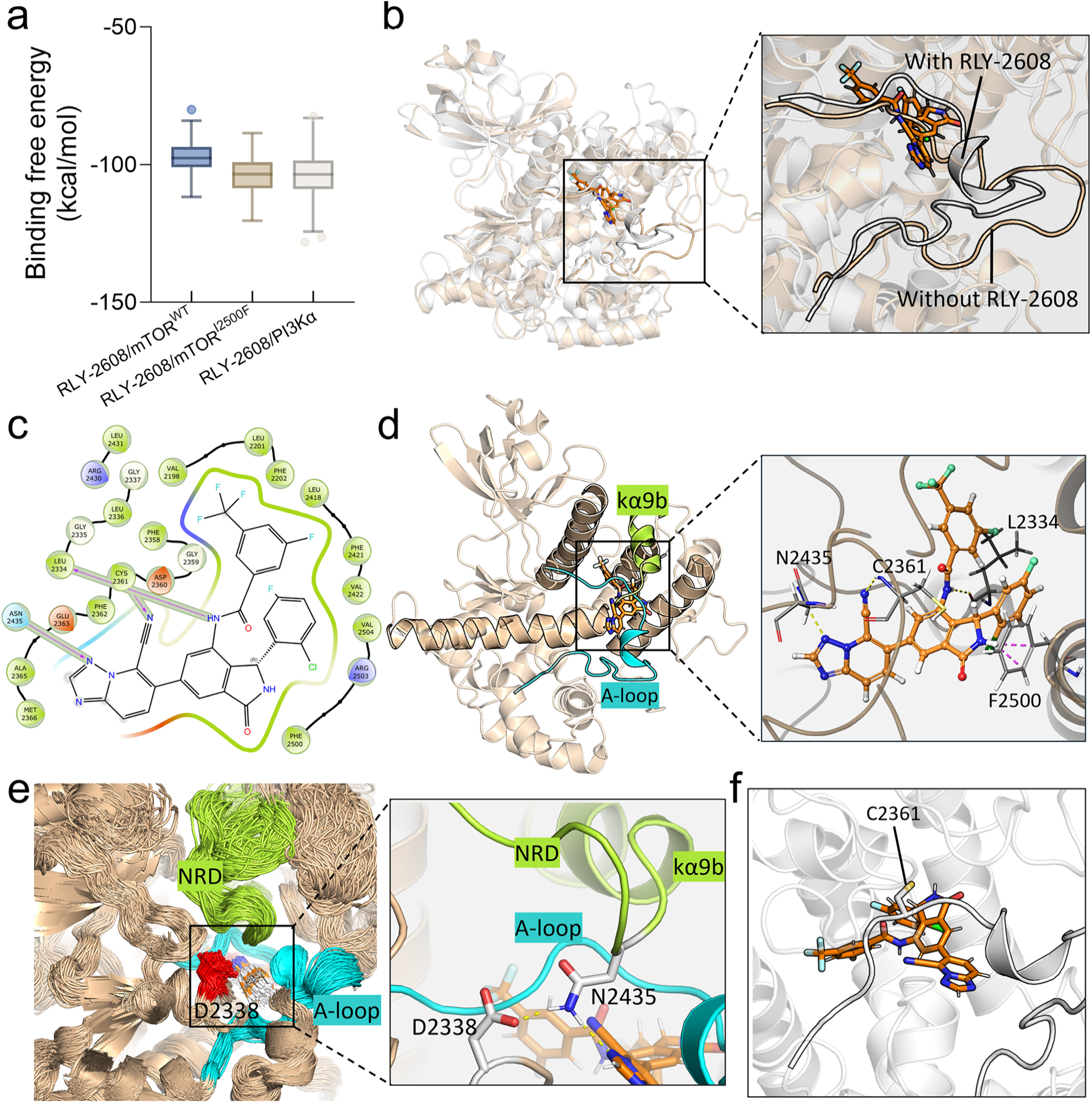
Interactions of RLY-2608 with I2500F mutant, WT mTOR, and PI3Kα. (a) Binding free energies between RLY-2608 and the I2500F mutant, WT mTOR, and PI3Kα. (b) Structural alignment of the I2500F mutant in the presence (white) and absence (wheat) of RLY-2608, illustrating the conformational changes in the A-loop induced by RLY-2608 binding. (c) A two-dimensional interaction map of RLY-2608 with the I2500F mutant mTOR. (d) H-bonds formed between residues in the I2500F mutant and RLY-2608. (e) The preferred orientation of D2338 in the I2500F mutant mTOR in the presence of RLY-2608. (f) A snapshot highlighting the C2361 residue in the A-loop, located near the newly discovered cryptic pocket.

Designing mutant-sensitive allosteric drugs for mTOR requires structure-based strategies to tailor the drug to effectively fit the mTOR cryptic pocket. We propose to screen or design allosteric inhibitors based on PI3Kα allosteric drugs to target this newly identified allosteric cryptic pocket in specific mutant mTORs. While WT mTOR typically maintains a stable α-helix packing, this mutant-selective pocket promotes potential allosteric drug specificity. Although other selected mutants did not exhibit a prominent allosteric pocket at this site within the simulation time, their ability to disrupt the α-packing leads us to believe that this pocket is likely to manifest on longer timescales. We expect that binding of such allosteric drugs to mutant mTORs can restore the α-helical structure of the kαAL-helix (or A-loop), stabilize the α-packing, maintain a closed catalytic cleft, and retain the orientation and position of active site residues less favorable for catalysis.

## DISCUSSION

Cancer is a genomic disease commonly driven by genetic and epigenetic alterations that disrupt the population balance between active and inactive proteins, thereby enhancing cell signaling for proliferation and growth. In this study, we analyzed cancer genomic databases to identify mTOR driver mutations. Using MD simulations, we showed that stable and precise FAT-KD interaction exerts an inhibitory effect on mTOR, consistent with prior experimental findings.^44, 62^ Deletion of the FAT domain can disrupt the packing formed by helices kαAL, kα3, kα9, kα9b, and kα10, typically leading to loss of helical components of the kαAL-helix, extension of the A-loop, and higher fluctuation of the kα9b-helix and the NRD, which partially eliminates the blockage of the catalytic cleft, and bring active site residues into closer proximity with ATP, favorable for catalysis. This explains why RHEB binding, which loosens the contact between the FAT and kinase domains (**Fig. S12**), can allosterically activate mTORC1, increasing the catalytic constant from 0.09 s□¹ to 2.9 s□¹.^44^

Additionally, we provide an atomic-level explanation for the activation of mutant mTORs. The selected cancer-associated mutations identified from the cancer genomic databases have relatively high frequencies of occurrence within the FAT and kinase domains of mTOR. These mutations include E1799K, T1977K, V2006F, S2215F, E2419K, L2427R, and I2500F. In our simulations, WT mTOR maintains a stable and inhibitory structure in the kinase domain. In contrast, all these mutations induce a cascade of activation events. Regardless of the mutation positions, these mutations can disrupt the packing structure formed by helices kαAL, kα3, kα9, kα9b, and kα10, aligning with prior hypotheses.^50, 51^ This disruption can be either a cause or a consequence, depending on the mutation positions. L2427R and I2500F, located within this packing structure, can directly disrupt it, likely followed by other activation events. Since the E1799K and E2419K mutations can loosen the contact between the FAT and kinase domains, they can mimic the effect of ΔFAT, finally resulting in the active site residues approaching or orienting toward ATP, which is favorable for catalysis. The T1977K, V2006F, and S2215F mutations can disrupt the interaction between the FAT^C-term^ and the KD^N-lobe^. This perturbation affects the FRB domain’s anchoring onto the N-lobe, increasing its dynamics. Consequently, there is significant movement of the FRB domain, resulting in an open catalytic cleft. This phenomenon was also observed in mTOR with NRD deleted in our previous work.^51^ Experimental results have demonstrated that NRD deletion increases mTOR activity by 3.5-fold in assays.^53^ Despite T1977K, V2006F, and S2215F being distant from the α-packing, they can also allosterically disrupt it, primarily due to mutual constraints between the FRB domain and NRD. The displacement of the FRB domain increases the mobility of the NRD, inducing outward motion and disorder in kα9b, further triggering conformational changes in the A-loop. Although we did not observe the opening of the catalytic cleft in the L2427R, I2500F, E1799K, and E2419K mutants, we believe that these mutations could potentially induce rotation of the FRB domain and opening of the catalytic cleft over a longer timescale due to their ability to disrupt (or disturb) the kαAL-kα3-kα9-kα9b-kα10 packing. The opening of the catalytic cleft facilitates substrate access to the bottom of the cleft for phosphorylation.

Our study showed that disruption of the α-packing could create a cryptic pocket within this region, which may require binding of an allosteric inhibitor to restore proper packing. Mutant-specific allosteric inhibitors, such as STX-478 and RLY-2608,^19, 20^ bind to an analogous site in PI3Kα as our newly identified allosteric pocket in mutant mTORs. Molecular docking and MD simulation results demonstrated that RLY-2608 can stably bind to the cryptic pocket identified in the mTOR I2500F mutant. This binding induces a conformational change in the A-loop, triggering an allosteric signal that regulates the orientation of active-site residues, making them unfavorable for catalysis.^75^ Structural modifications of the drug could optimize its specificity. Especially, this cryptic PI3K-analog pocket could be a target for other specific allosteric inhibitors for mutant mTORs, expanding allosteric designs. We propose to screen for, or design, allosteric inhibitors based on PI3Kα allosteric drugs to target this novel cryptic pocket in specific mutant mTORs. Since WT mTOR typically maintains a stable α-packing, this pocket is less likely to form in WT mTOR, making the allosteric drugs selective for mutant forms.

### Conclusions

In conclusion, mTOR driver mutations enhance mTOR signaling in cell proliferation, and influence the response of the pathway to nutrient starvation, thereby benefiting cell growth. Structurally, the stable and precise interaction between the FAT and kinase domains inhibits its activity. Mutations at the FAT-KD interface (e.g. E1799K, E2419K) and RHEB binding can weaken this interaction, triggering conformational changes in the kinase domain. This results in the realignment of active site residues that promote catalysis. Mutations such as T1977K, V2006F, and S2215F, which disrupt the anchoring of the FAT^C-term^ to the KD^N-lobe^, can cause movement of the FRB domain and subsequent opening of the catalytic cleft. This increases the accessibility of the substrate to ATP. Across various mutation sites, all selected mTOR mutations disrupt the packing structure formed by the kαAL, kα3, kα9, kα9b, and kα10 helices. This disruption can involve loss of helical components in kαAL or kα9b, extension of the A-loop, or outward movement of kα9b and NRD, potentially releasing constraints on the active site. Such disturbances can create a cryptic pocket, likely necessitating an allosteric inhibitor to restore the packing structure to inhibit mutant mTORs. Although the opening of the catalytic cleft or the orientation of the active site towards ATP was not observed within the simulation timeframe for certain mutant mTORs, we believe that these activation events may occur over a longer simulation timescale. Molecular docking and MD simulations of the PI3Kα allosteric inhibitor RLY-2608 with the I2500F mutant and WT mTOR inspire new, RLY-2608 informed, allosteric inhibitors promising selective targeting of the mutant mTOR over WT.

Our findings decipher the activation mechanisms of mTOR variants, discovering a PI3K-like cryptic pocket. This, coupled with *learning* from the allosteric inhibitors of the evolutionary and mechanistically similar PI3Kα, epitomize that mechanistic knowledge can productively expand innovative allosteric inhibitor designs and strategies. Identifying which mutants respond effectively to specific drugs could improve patient selection and enable more targeted treatment strategies.

## Materials and Methods

### Analysis of cancer datasets

We used the TCGA and GENIE cancer databases to identify the significant mTOR mutations associated with cancer. The MC3 MAF file from the TCGA dataset and the mutation extended file of the GENIE cohort (v13.1-public) were downloaded from https://gdc.cancer.gov and https://www.synapse.org, respectively. We focused on missense mutations in mTOR, excluding deletion-insertion (delins) variations. To minimize the inclusion of incidental mutations, we removed data with a variant allele frequency (VAF) ≤ 0.125 and an occurrence frequency ≤ 3. Data processing and statistical analysis were conducted using code of Python 3.10, and mutation frequency plots were generated using the ProteinPaint tool (https://proteinpaint.stjude.org).^76^

### Construction of simulation systems

In this study, we modeled WT mTOR, ΔFAT, and seven mutant mTOR systems: E1799K, T1977K, V2006F, S2215F, E2419K, L2427R, and I2500F for simulations (**Table S1**). All systems were constructed based on the crystal structure of N-terminal truncated mTOR complexed with mLST8 (PDB ID: 4JSP). Missing regions in the crystal structure were modeled using the AlphaFold2 package.^77, 78^ Given the active site restriction mechanism, in which the kα9b-helix and NRD block the mTOR active site, we manually adjusted the NRD to position it closer to the active site. The ΔFAT system was generated by deleting a large portion of the FAT domain (residues 1385-1934), while retaining the fα35-helix (**Fig. S2**). The E1799K, T1977K, V2006F, S2215F, E2419K, L2427R, I2500F mutant systems were constructed by substituting residues E1799, T1977, V2006, S2215, E2419, L2427, and I2500 with residues Lys, Lys, Phe, Phe, Lys, Arg, and Phe, respectively. An ATP molecule and two magnesium ions (Mg^2+^) were placed in the ATP-binding pocket of mTOR in all complexes. Each system was solvated using the TIP3P solvent model, and Na^+^ and Cl^−^ were added to the systems to neutralize and achieve a physiological salt concentration of ∼0.15 mol L^−1^.

### MD simulation protocol

The all-atom MD simulations were conducted for all systems, employing the NAMD 2.14 package^79^ with the CHARMM^80^ all-atom additive force field (version C36m).^81, 82^ The simulation protocol in this work closely follows that used in our previous works.^4, 5, 83, 84, 85^ Prior to production runs, we performed multiple cycles of minimization and dynamics to eliminate any bad contacts between atoms within the systems. In the production runs, each system underwent 3 independent MD simulations within the NPT ensemble, maintaining 3D periodic boundary conditions. Each independent simulation was run for 3 μs. Notably, the results derived from these parallel trajectories consistently exhibited similarities and were considered comparable. Throughout the simulations, we maintained a pressure of 1 atm using the Langevin piston control algorithm, while the temperature was preserved at 310 K using the Langevin thermostat method with a damping coefficient of 1 ps^-1^. Covalent bonds, including hydrogen atoms, were constrained using the SHAKE algorithm, and a time-step of 2 fs was employed. We utilized the particle mesh Ewald method and switching functions for the computation of long-range electrostatic and short-range van der Waals (vdW) potential energies, respectively. Subsequently, the analysis was carried out using the FORTRAN script within the CHARMM package (version c45b1), the TCL script within the VMD package, and the Python script within the MDAnalysis package.

### Molecular docking, modeling, and MD simulation for RLY-2608 interaction with WT mTOR, mTOR I2500F mutant, and PI3K**α**

We first aligned the final structures of the kinase domain of WT mTOR and the I2500F mutant with the complex structure of PI3Kα and RLY-2608 (PDB ID: 8TS7). Induced Fit Docking (IFD) was then performed using Maestro^86^ to dock RLY-2608 to WT mTOR, the I2500F mutant, and PI3Kα, and we selected the configurations with the best IFDScore for further MD simulations. The complex systems—RLY-2608/mTOR^WT^, RLY-2608/mTOR^I2500F^, and RLY-2608/PI3Kα (**Table S2**)—were parameterized using the OPLS4 force field and solvated with the TIP3P model. Na and Cl ions were added to neutralize the systems and achieve a physiological salt concentration of approximately 0.15 mol/L. Each system underwent 10 ps of energy minimization prior to the MD simulations. We executed three parallel MD trajectories for each system using Desmond,^87^ with each trajectory running for 500 ns under the NPT ensemble. The temperature and pressure were maintained at 310 K and 1 atm using the Nose-Hoover Chain and Martyna-Tobias-Klein algorithms, respectively. The cutoff for Coulombic interactions was set to 10.0 Å, and a time step of 2 fs was employed. The binding free energies between the ligand and receptor were calculated using the MM-GBSA module in Maestro.

## Author Contributions

YL, WZ, HJ, and RN conceived and designed the study. YL conducted the simulations. YL performed the bioinformatic analysis, analyzed the simulation results, and drafted the manuscript. YL, WZ, HJ, and RN validated, reviewed, and edited manuscript. RN supervised the project.

## Declaration of interests

None

## Acknowledgments

This project has been funded in whole or in part with federal funds from the National Cancer Institute, National Institutes of Health, under contract HHSN261201500003I. The content of this publication does not necessarily reflect the views or policies of the Department of Health and Human Services, nor does mention of trade names, commercial products, or organizations imply endorsement by the US Government. This research was supported [in part] by the Intramural Research Program of the NIH, National Cancer Institute, CCR. The calculations had been performed using the high-performance computational facilities of the Biowulf PC/Linux cluster at the National Institutes of Health, Bethesda, MD (https://hpc.nih.gov/).

The authors declare no potential conflicts of interest.

## Notes

### Competing Interest Statement

The authors have declared no competing interest.

