## Supporting information of figures and tables for "mTOR variants activation discovers PI3K-like cryptic pocket, expanding allosteric, mutant-selective inhibitor designs"

The authors declare no potential conflicts of interest.

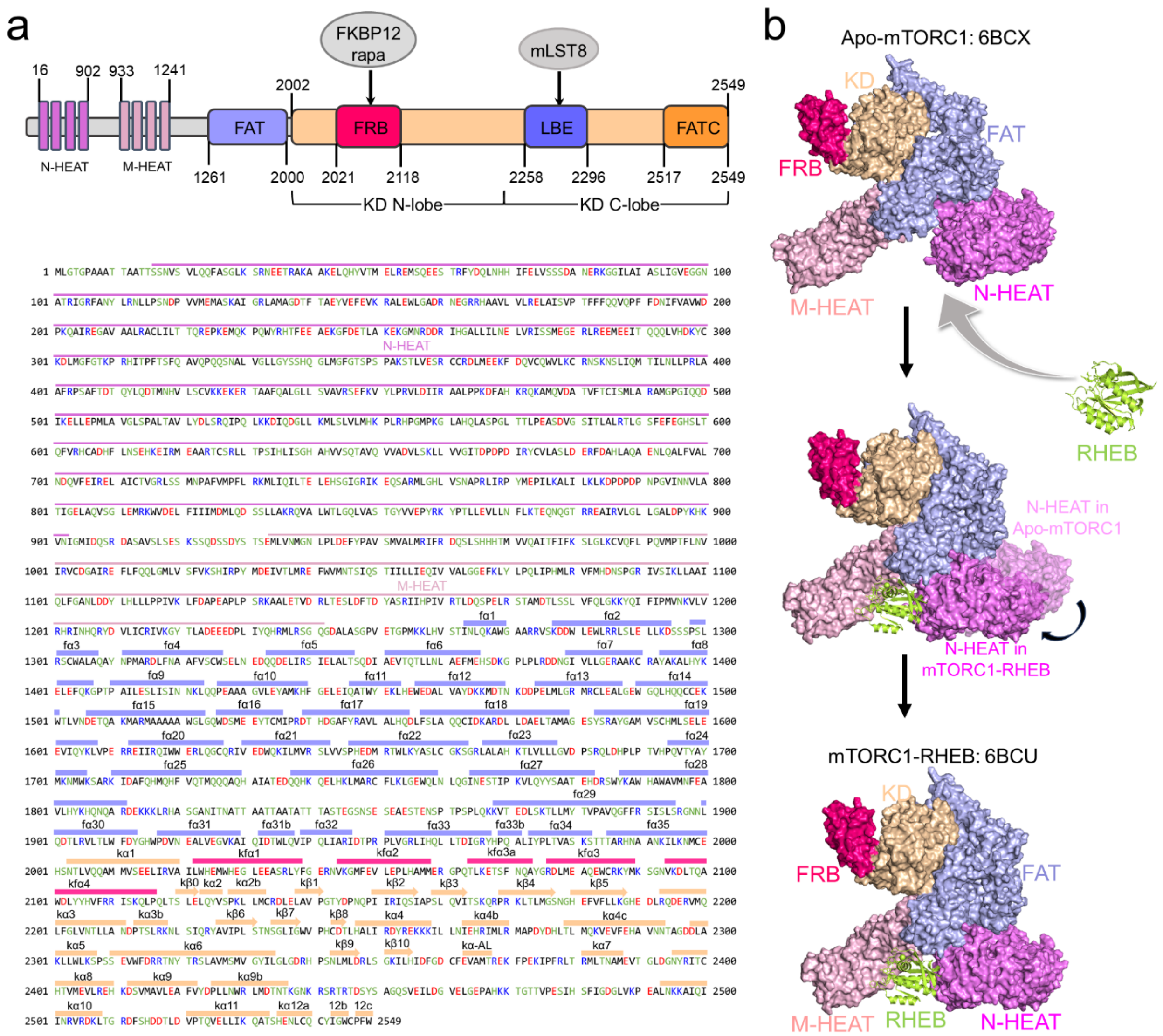


**Fig. S1**. **Domain structure and sequence of mTOR.** (a) mTOR comprises two N-terminal HEAT regions: N-HEAT (residues 16-902) and M-HEAT (residues 933-1241), followed by the FAT domain (residues 1261-2000) and the kinase domain (residues 2002-2549). Within the kinase domain, three inserted domains exist: the N-lobe inserted FRB domain (residues 2021-2118) responsible for rapamycin-FKBP12 complex binding, and the C-lobe inserted LBE (residues 2258-2296) and FATC (residues 2517-2549) domains. In both mTORC1 and mTORC2, the LBE domain binds mLST8. (b) Allosteric activation mechanism of mTORC1 induced by RHEB binding. In the PI3K/AKT/mTOR pathway, RHEB acts as the physiological allosteric activator of mTORC1.[1, 2] Cryo-EM structures of the mTORC1–RHEB complex (PDB ID: 6BCU) and apo-mTORC1 (PDB: 6BCX) reveal that RHEB binding to N-HEAT of mTOR causes a conformational change, pulling N-HEAT closer to M-HEAT and twisting the middle region of the FAT domain. This changes the relative orientation of active site residues and brings ATP phosphate groups closer to critical catalytic residues.

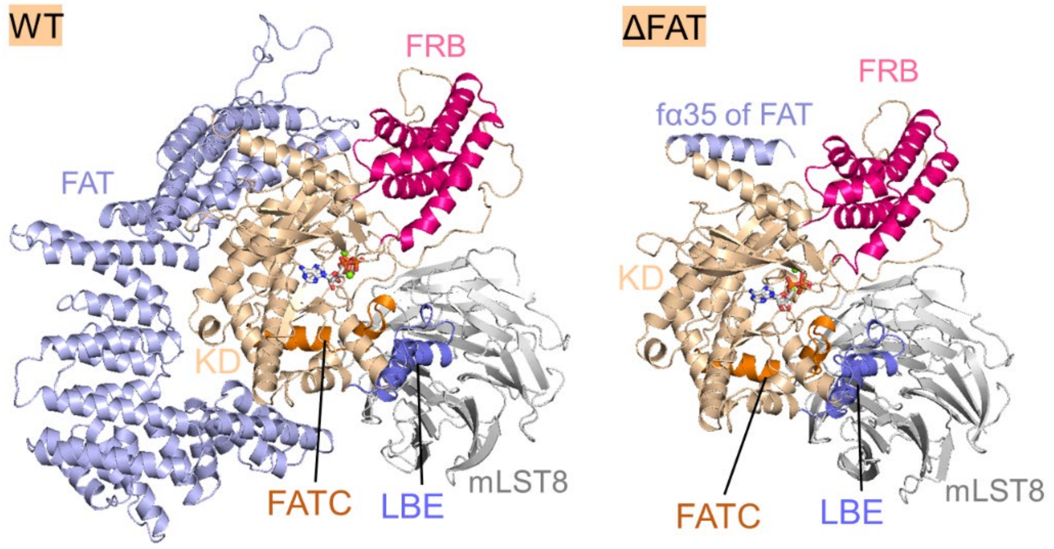


**Fig. S2. Initial configurations of mTOR model systems for simulations.** Initial configurations of the WT (*left panel*) and ΔFAT (*right panel*) systems were constructed based on the crystal structure (PDB ID: 4JSP).

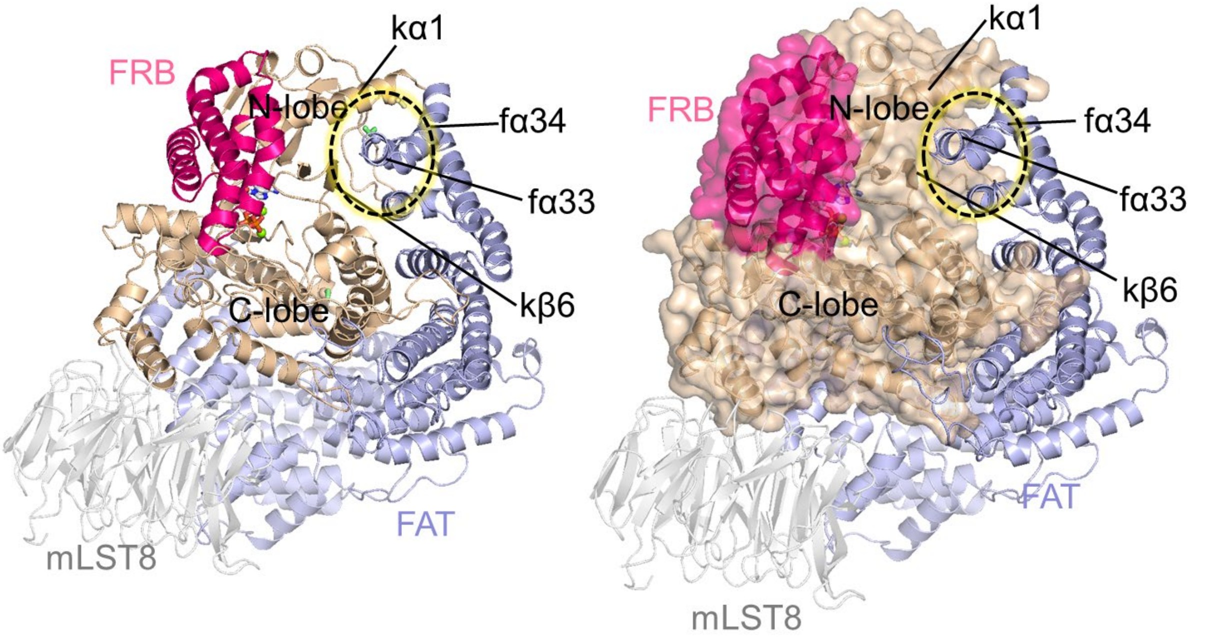


**Fig. S3. Snapshots illustrating the anchoring of the C-terminus of the FAT domain onto the N-lobe of the kinase domain of mTOR.** The kinase domain of mTOR including the FRB domain is shown as a cartoon or surface representation. The FAT domain and mSLT8 are shown as cartoons. The dotted circle indicates the anchorage area.

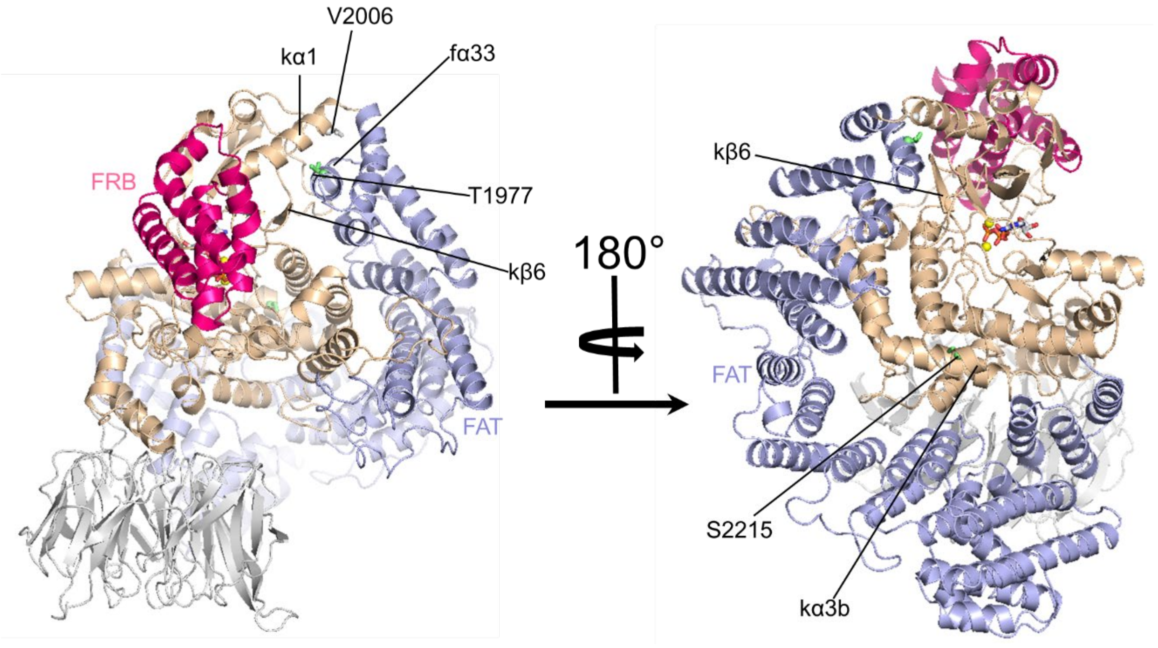


**Fig. S4**. **Mapping of the T1977K, V2006F and S2215F mutations on the mTOR structure.** Snapshot showing the positions of residues T1977, V2006, and S2215 within the fα33-helix, kα1-helix, and kα3b-helix, respectively.

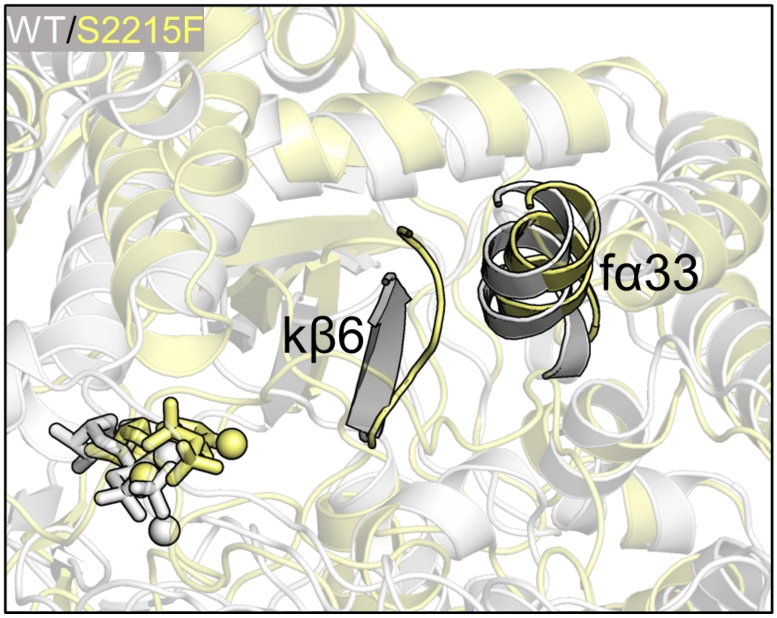


**Fig. S5**. **Structural alignment of the S2215F mutant with respect to WT mTOR.** Carttons highlighting the conformational change in the kβ6-strand induced by the S2215F mutation.

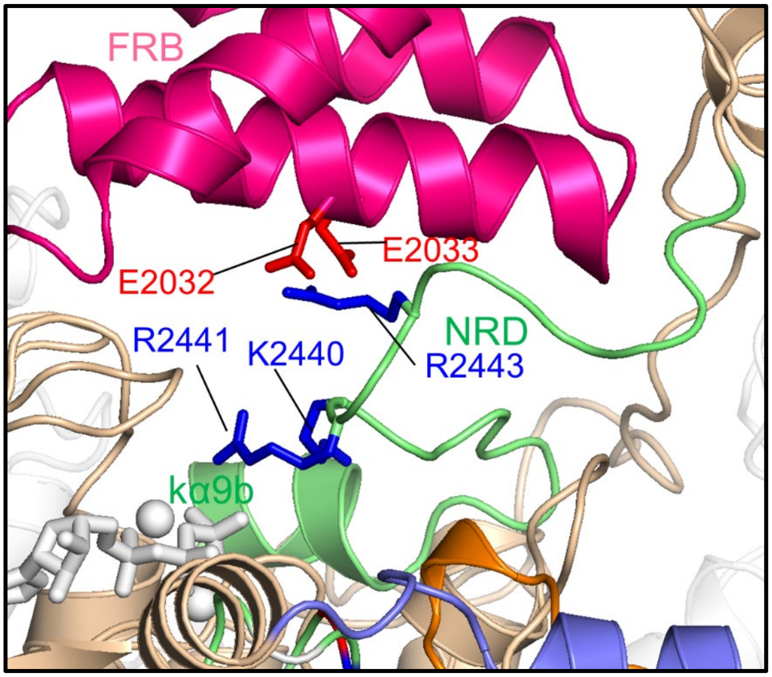


**Fig. S6**. **Constraint of NRD in the in the catalytic cleft.** Snapshot depicting electrostatic or salt bridge interactions between positively charged residues (K2440, R2441, and R2443) in the NRD and negatively charged residues (E2032 and E2033) in the FRB domain.

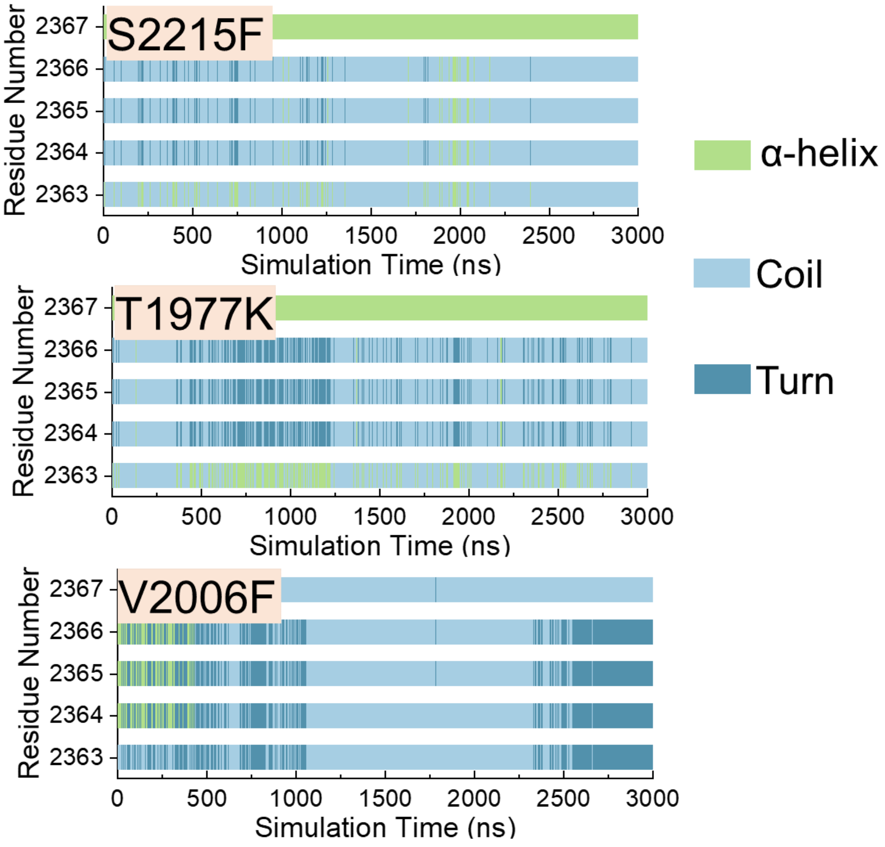


**Fig. S7. Conformational change of the kαAL-helix for mutant mTORs.** Time evolution of the secondary structure of the kαAL-helix for the S2215F, T1977K, and V2006F mutants.

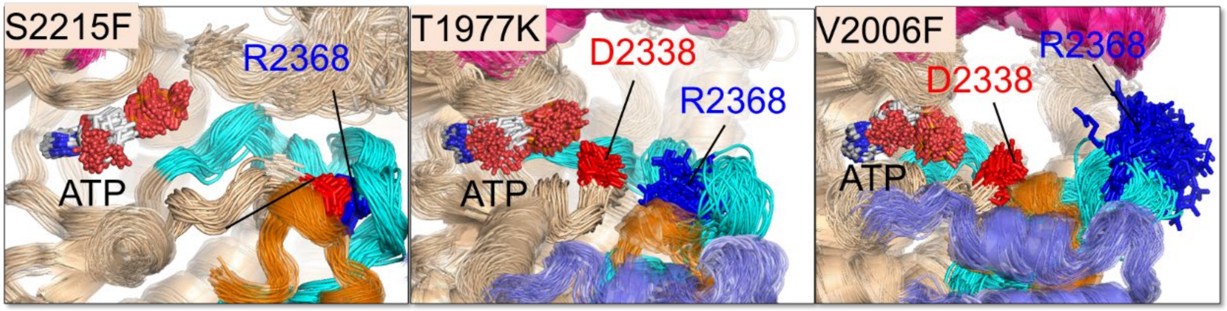


**Fig. S8**. **Clustered snapshots highlighting the catalytic residue D2338.** Relative orientations of D2338 in the S2215F, T1977K, and V2006F mutants over the last 1 μs trajectories.

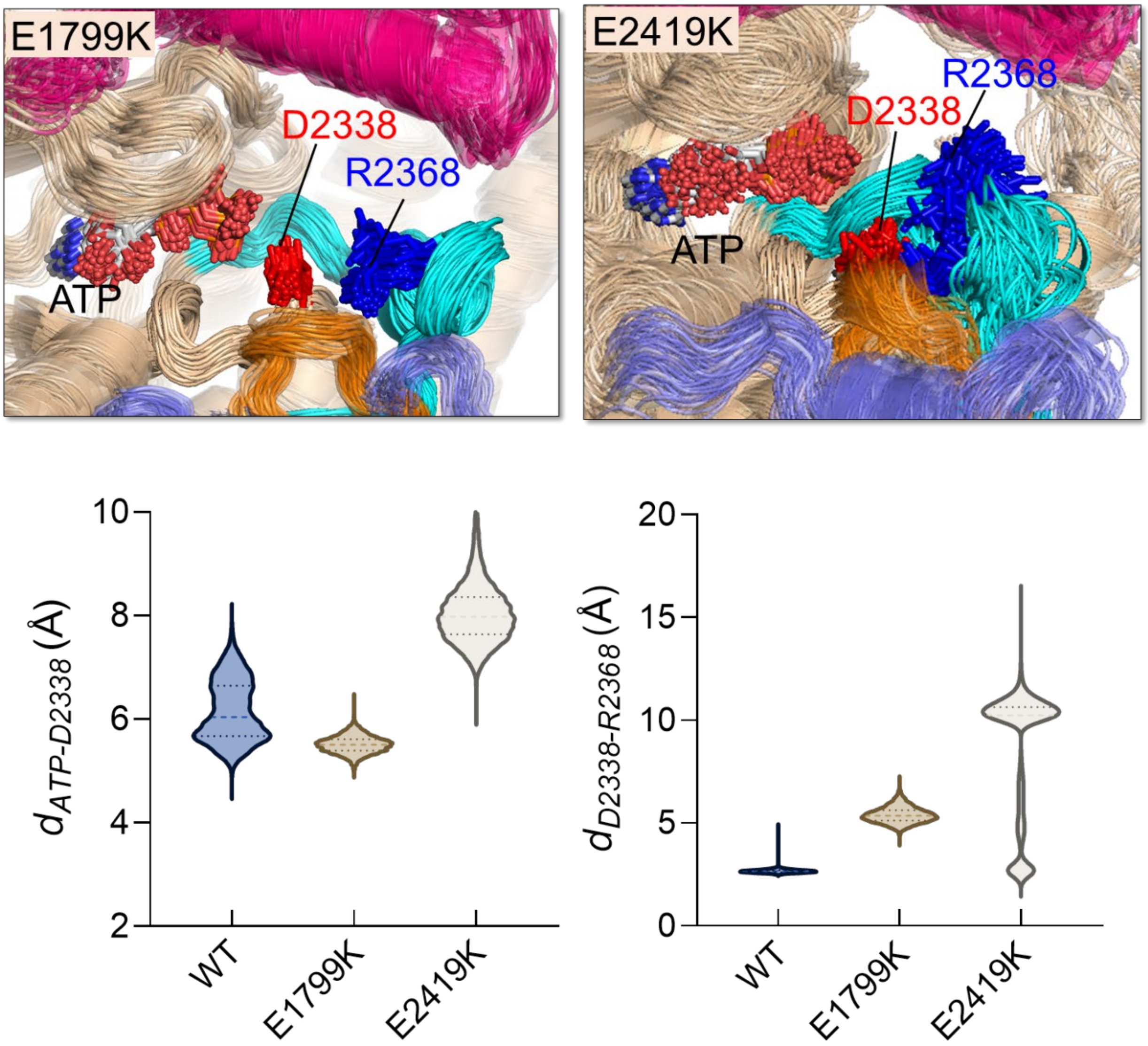


**Fig. S9. Conformational ensembles of the A-loop and the distances between key atom pairs for the E1799K and E2419K mutant systems.** Clustered snapshots highlighting R2368 in the A-loop and D2338 in the catalytic loop (Cat-loop) (*top panels*). The distances between ATP:PG and D2338:OD2 ($d_{ATP–D2338}$) and between D2338:OD2 and R2368:NH1 ($d_{D2338–R2368}$) (*bottom panels*). Conformations and data were extracted from the last 1 µs of trajectories.

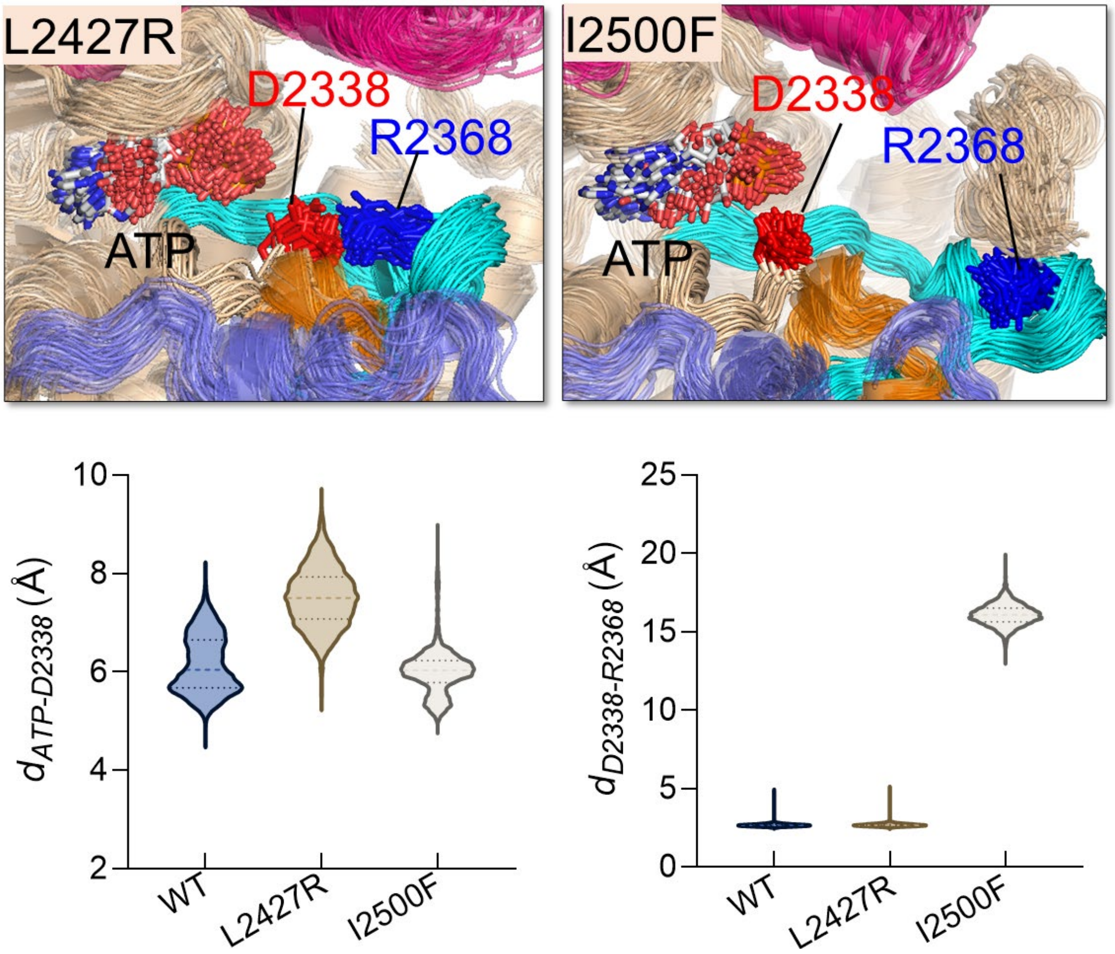


**Fig. S10. Conformational ensembles of the A-loop and the distances between key atom pairs for the l2427R and I2500F mutant systems.** Clustered snapshots highlighting R2368 in the A-loop and D2338 in the catalytic loop (Cat-loop) (*top panels*). The distances between ATP:PG and D2338:OD2 ($d_{ATP–D2338}$) and between D2338:OD2 and R2368:NH1 ($d_{D2338–R2368}$) (*bottom panels*). Conformations and data were extracted from the last 1 μs of trajectories.

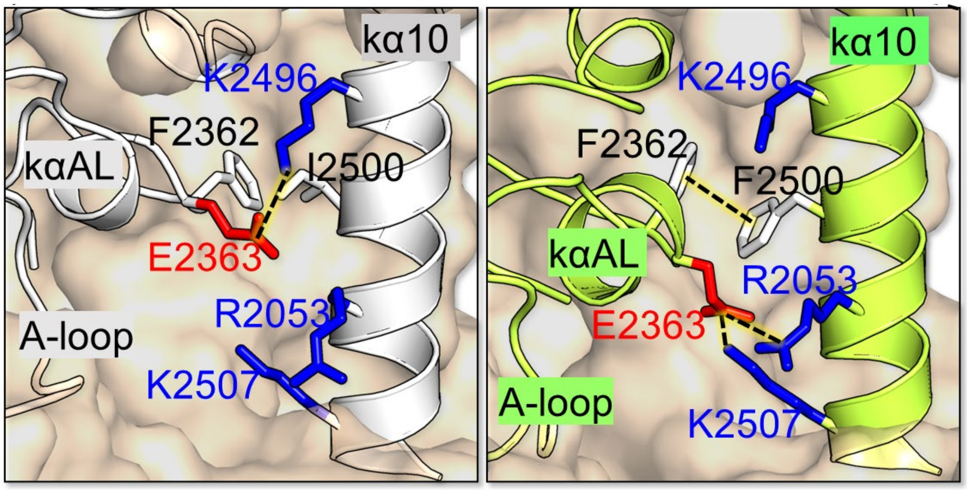


**Fig. S11. Formation of π-π stacking in the I2500F mutant mTOR.** Snapshots showing the interaction between residues in the kα10-helix and residues in the A-loop for the WT (*left*) and I2500F (*right*) systems. The π-π stacking between F2362 and F2500 is responsible for the formation of the cryptic allosteric pocket.

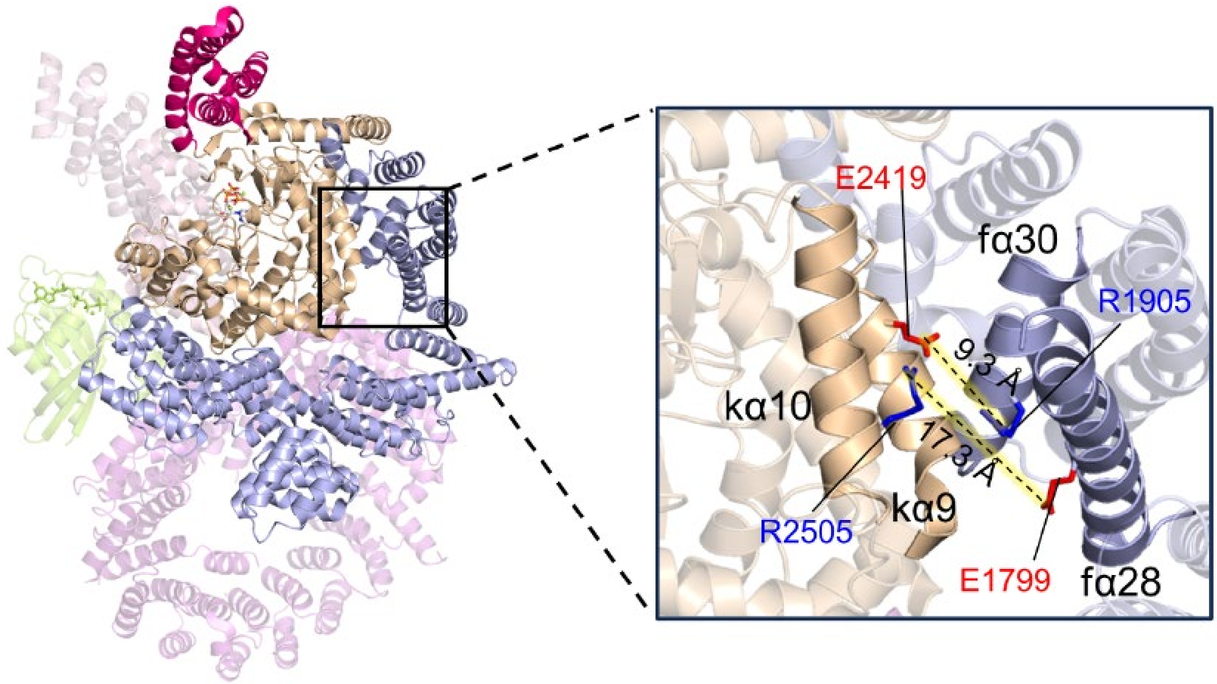


**Fig. S12. Interaction of the FAT domain with the kinase domain.** Cryo-EM structure of the mTORC1–RHEB complex (PDB ID: 6BCU) showing disruption of the interface between the FAT domain and the α-packing of the kinase domain induced by binding of RHEB to the N-HEAT domain of mTOR in mTORC1. In our simulation, the E2419-R1905 and E1799-R2505 salt bridges can be maintained in WT mTOR. In the cryo-EM structures, the distances between these paired residues exceed the 4.0 Å cutoff for salt bridges, indicating disruption of the interface.

**Table S1.** Details of simulation systems.

| System name | Components | mTOR mutation | Mutation motif | No. of parallel trajectories | Simulation time |
|---|---|---|---|---|---|
| WT | mTOR, mLST8, ATP, 2 $Mg^{2+}$ | None | / | 3 | 3 μs |
| ΔFAT | ΔFAT-mTOR, mLST8, ATP, 2 $Mg^{2+}$ | Deletion of FAT domain | / | 3 | 3 μs |
| E1799K | mTOR, mLST8, ATP, 2 $Mg^{2+}$ | E1799K | fα28 | 3 | 3 μs |
| T1977K | mTOR, mLST8, ATP, 2 $Mg^{2+}$ | T1977K | fα34 | 3 | 3 μs |
| V2006F | mTOR, mLST8, ATP, 2 $Mg^{2+}$ | V2006F | kα1 | 3 | 3 μs |
| S2215F | mTOR, mLST8, ATP, 2 $Mg^{2+}$ | S2215F | kα3b | 3 | 3 μs |
| E2419K | mTOR, mLST8, ATP, 2 $Mg^{2+}$ | E2419K | kα9 | 3 | 3 μs |
| L2427R | mTOR, mLST8, ATP, 2 $Mg^{2+}$ | L2427R | kα9b | 3 | 3 μs |
| I2500F | mTOR, mLST8, ATP, 2 $Mg^{2+}$ | I2500F | kα10 | 3 | 3 μs |

Note: Here, mTOR refers to the N-terminal truncated mTOR (residues 1382-2549).

**Table S2.** Details of simulation systems for RLY-2608 interaction with WT mTOR, I2500F mutant, and PI3Kα.

| System name | Components | Coordinates of receptor | No. of parallel trajectories | Simulation time |
|---|---|---|---|---|
| RLY-2608/mTOR$^{WT}$ | Kinase domain of WT mTOR, RLY2608 | Final frame of the WT system | 3 | 500 ns |
| RLY-2608/mTOR$^{I2500F}$ | Kinase domain of I2500F mutant mTOR, RLY2608 | Final frame of the I2500F system | 3 | 500 ns |
| RLY-2608/PI3Kα | Kinase domain of PI3Kα, RLY2608 | PDB ID: 8TS8 | 3 | 500 ns |
